# Inhibition of Ca_V_1.4 channels by Ca_V_3 channel antagonists ML218 and Z944

**DOI:** 10.1101/2025.08.30.673144

**Authors:** Jinglang Sun, Juan de la Rosa Vázquez, Adriana Hernández-González, Vladimir Yarov Yarovoy, Amy Lee

**Author notes:** Corresponding author: Amy Lee Dept. of Neuroscience The University of Texas at Austin 100 E. 24^th^ St. Austin TX 78712, USA.

## Abstract

Among the three classes of voltage-gated Ca^2+^ channels (Ca_v_1, Ca_v_2, Ca_v_3), Ca_v_3 T-type channels are drug targets for disorders including epilepsy and pain. Antagonists such as Z944 and ML218 are highly selective for Ca_v_3 compared to the Ca_v_1.2 L-type channel but whether they have additional activity on other Ca_v_1 subtypes is unknown. Here, we investigated the effects of Z944 and ML218 on the Ca_v_1.4 channel which regulates neurotransmitter release from retinal photoreceptors. In HEK293T cells transfected with Ca_v_1.4 and the auxiliary β_2×13_ and α_2_δ-4 subunits, Z944 and ML218 inhibited Ca^2+^ currents with IC_50_ values of ∼30 µM and 2 µM, respectively. Structure-based modeling combined with functional studies revealed the importance of a cluster of methionine residues, particularly M1004, within the DHP binding site for the effects of ML218. Compared to mutation of a conserved threonine (T1007) that is required for DHP sensitivity of Ca_v_1 channels, mutation of M1004 had a 10-fold greater impact in diminishing the potency of ML218. Ca_v_1.2 was significantly less sensitive to ML218 inhibition (IC_50_∼ 37 µM) than Ca_v_1.4, which could not be attributed to a valine in place of M1004 in Ca_v_1.2. We conclude that ML218 and Z944 are dual Ca_v_1/Ca_v_3 modulators of Ca_V_1.4 and should be used with caution when dissecting the contributions of Ca_V_3 channels in tissues where Ca_v_1.4 is expressed.

## INTRODUCTION

Ca_V_3 T-type Ca^2+^ channels (Ca_V_3.1, Ca_V_3.2, Ca_V_3.3) are important regulators of cellular excitability in a wide array of tissues. Compared to Ca_V_1 and Ca_V_2 channels, Ca_V_3 channels activate and inactivate at hyperpolarized voltages which enable their contribution near the resting potential of many neurons (Perez-Reyes, 2003). Modulation of Ca_V_3 channels leads to specific patterns of neuronal activity including low-threshold Ca^2+^ spikes (Llinas and Yarom, 1981), burst firing (Huguenard and Prince, 1992), and rhythmic oscillations (Williams et al., 1997; Hughes et al., 2002). Loss- or gain-of function of Ca_V_3 channels is linked to a variety of disorders including epilepsy, chronic pain, autism spectrum disorder, and primary aldosteronism (Weiss and Zamponi, 2020). Given their physiological importance, Ca_V_3 channels have been the subject of intense research as major drug targets.

Ethosuximide is one of the first Ca_V_3 blockers to be used clinically for the treatment of absence epilepsy but it can have actions on other ion channels (Shalomov et al., 2025) as well as adverse side effects (Goren and Onat, 2007). After the molecular cloning of the Ca_V_3 channels, Z944 and ML218 were developed as more selective blockers of these channels. Z944 is a piperazine derivative with sub-micromolar affinity for the three Ca_V_3 subtypes that is ∼70-100 times higher than that for Ca_V_2.2 and Ca_V_1.2 (Tringham et al., 2012) (Supp. Fig.1A). Based on strong pre-clinical evidence (Lee, 2014; Harding et al., 2021; Scott et al., 2022; Matthews et al., 2023), Z944 is currently in phase II and phase III clinical trials for pain and essential tremor, respectively (Weiss and Zamponi, 2019; Giroux et al., 2024). Following a high throughput screening assay for small-molecules with activity on Ca_V_3 channels, ML218 was derived from one of the hits using a scaffold-hopping approach (Supp. Fig.1A). Like Z944, ML218 has high affinity for Ca_V_3 channels (IC_50_ < 500 nM) and significantly inhibits Ca_V_3 current and rebound burst firing in subthalamic nucleus neurons (Xiang et al., 2011). However, ML218 failed to have similar effects in Parkinsonian monkeys (Galvan et al., 2016), despite having anti-Parkinsonian effects in a rat model of catalepsy (Xiang et al., 2011). Due to their potent inhibition of Ca_V_3 channels, Z944 and ML218 have been extensively used to assay the contributions of Ca_V_3 channels to neuronal excitability, physiology, and behavior (Matschke et al., 2015; Li et al., 2017; Roebuck et al., 2018; Davison et al., 2022; Baggio et al., 2024).

While Z944 and ML218 have significantly weaker affinity for Ca_V_1.2 than any of the Ca_V_3 subtypes (Xiang et al., 2011; Tringham et al., 2012), recent evidence indicates that when used in the micromolar range, these drugs can have actions on presynaptic Ca_V_ currents in cone photoreceptors of the mouse retina (Davison et al., 2022; Maddox et al., 2024). Based on a wealth of evidence from human genetics and animal studies, the channel mediating these currents is Ca_V_1.4 (reviewed in (Williams et al., 2022)). Like dihydropyridine (DHP) Ca_V_1 agonists (e.g., BayK 8644 and FPL 64176), Z944 was found to potentiate Ca_V_1.4 currents in cones, causing a hyperpolarizing shift in channel activation (Davison et al., 2022). ML218 had similar effects in cones of mouse and ground squirrel retina, but also suppressed peak current amplitudes (Maddox et al., 2024). The dual modulatory effects of ML218 and Z944 are reminiscent of the opposite effects of optical isomers of some dihydropyridines on Ca_V_1 L-type Ca^2+^ channels (Schramm et al., 1983a; Franckowiak et al., 1985).

Here, we investigated the molecular basis of these effects of Z944 and ML218 in whole-cell patch-clamp recordings of HEK293T cells transfected with Ca_V_1.4 and auxiliary β_2×13_ and α_2_δ-4 subunits. We report that low micromolar concentrations of ML218 and Z944 significantly inhibit Ca_V_1.4 current density and that ML218 has minor effects as an agonist. Structure-based modeling and mutational analyses support a mechanism whereby these drugs interact in the DHP binding site of Ca_V_1.4.

## MATERIALS AND METHODS

### cDNAs and molecular cloning

The cDNAs for human Ca_V_3.2 (GenBank: AF051946), human Ca_V_1.4 (GenBank: AF201304), rabbit Ca_V_1.2a (GenBank: NP_001129994), human β_2×13_ (GenBank: AF465485), human α_2_δ-4 (GenBank: NM_172364), and enhanced GFP in pEGFP-C1 were used for co-transfection experiments. The Q5 Site-Directed Mutagenesis Kit (New England Biolabs) was used to generate the Ca_v_1.4 T1007 cDNA. The HiFi DNA Assembly Cloning System (New England Biolabs) was used to generate Ca_V_1.4 mutants with substitutions of alanine for methionine at positions 1004, 1129, or 1426 (M1004A, M1129A, M1426A) and the Ca_V_1.2 mutant with substitution of methionine for valine at position 1063 (V1063M). Primer sequences used for the corresponding mutagenesis reactions are listed in Table 1.

**Table 1.**
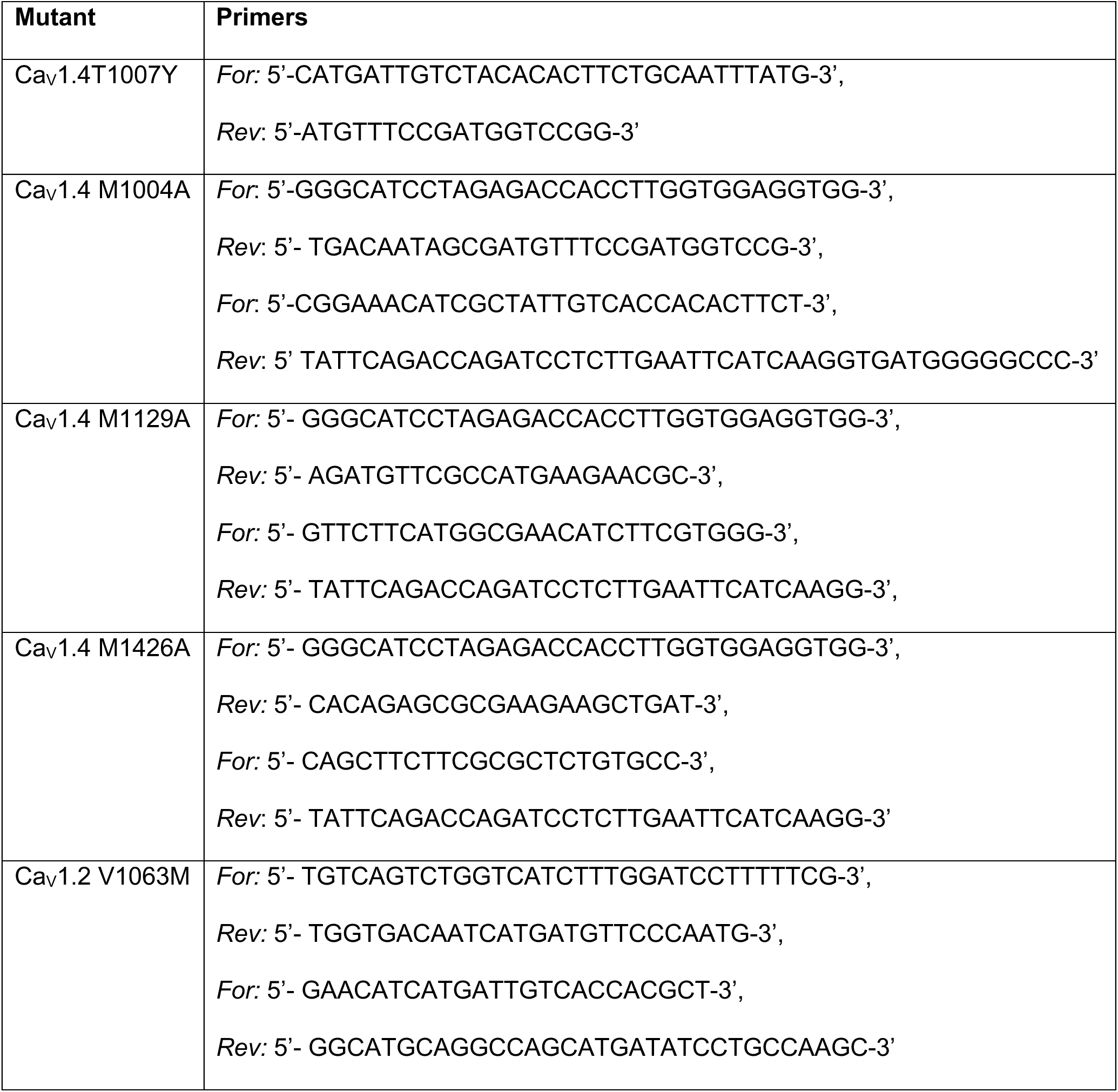
List of primers used for generating Ca_v_ mutant cDNAs.

### Cell culture and transfection

Human embryonic kidney 293 cells expressing SV40 T-antigen (HEK293T) were obtained from the American Type Cell Culture Collection (ATCC CRL-11268) and cultured in Dulbecco’s Modified Eagle’s Medium (Sigma-Aldrich) supplemented with 10% fetal bovine serum (VWR) and 1% penicillin–streptomycin (Sigma-Aldrich), at 37 °C in a humidified atmosphere of 5% CO₂. Cells cultured to 70-80% confluence in 35 mm petri dishes and were co-transfected with cDNAs encoding Ca_V_3.2, Ca_V_1.4, Ca_V_1.2, or a channel mutant (1.6 μg), β_2×13_ (0.6 μg), α_2_δ-4 (0.6 μg), and eGFP (0.15 μg) using Fugene 6 transfection reagent (Promega) according to the manufacturer’s protocol. Cells treated with transfection mixture were incubated for 24-48 hours at 37 °C prior to being dissociated via TrypLE Express solution (ThermoFisher) and plated at low density for single cell electrophysiological recordings.

### Drugs

Stock solutions (10 mM) of ML218 hydrochloride (Tocris, Cat. No. 4507) and Z944 (Tocris, Cat. No. 6367) were dissolved in dimethyl sulfoxide (DMSO) and H_2_O, respectively and stored at –20 °C. Serial dilutions of ML218 or Z944 were made from fresh aliquots of stock solutions by diluting in external recording solution to the indicated concentrations. The vehicle solution for experiments involving ML218 contained up to 1% DMSO.

### Electrophysiological Recordings

Whole-cell patch-clamp recordings were performed at room temperature between 36 and 72 hours post-transfection using an EPC10 amplifier or an EPC-8 amplifier and InstruTECH LIH 8+8 data acquisition system with PatchMaster software (HEKA Elektronik). External recording solution contained (in mM): 140 Tris, 1 MgCl₂, and 10 CaCl₂ or 20 CaCl₂. Internal recording solution contained (in mM): 140 N-methyl-D-glucamine, 10 HEPES, 10 EGTA, 2 MgCl₂, and 2 Mg-ATP. The pH of both solutions was adjusted to 7.3 with methanesulfonic acid. Electrodes were made from thin-walled borosilicate glass capillaries (World Precision Instruments) using a P-1000 Flaming/Brown micropipette puller (Sutter Instrument). The electrodes had resistances of 4–8 MΩ in the bath solution. Series resistance was compensated to 50–70%. Leak currents were subtracted using a P/–4 protocol. Recording files were analyzed using Igor Pro (Wavemetrics) and MATLAB (MathWorks) scripts written to extract peak current values, and analyzed data are presented as mean ± SEM.

Drug- or vehicle-containing solutions were delivered via a pressurized multichannel perfusion system connected to a perfusion pencil with a 250-μm perfusion tip (AutoMate Scientific). For dose-response relationships, currents were elicited at a frequency of 0.1 Hz and the maximal inhibition was calculated when the current amplitude reached a steady state following drug application (Suppl. Fig.2).

**Figure 1.**
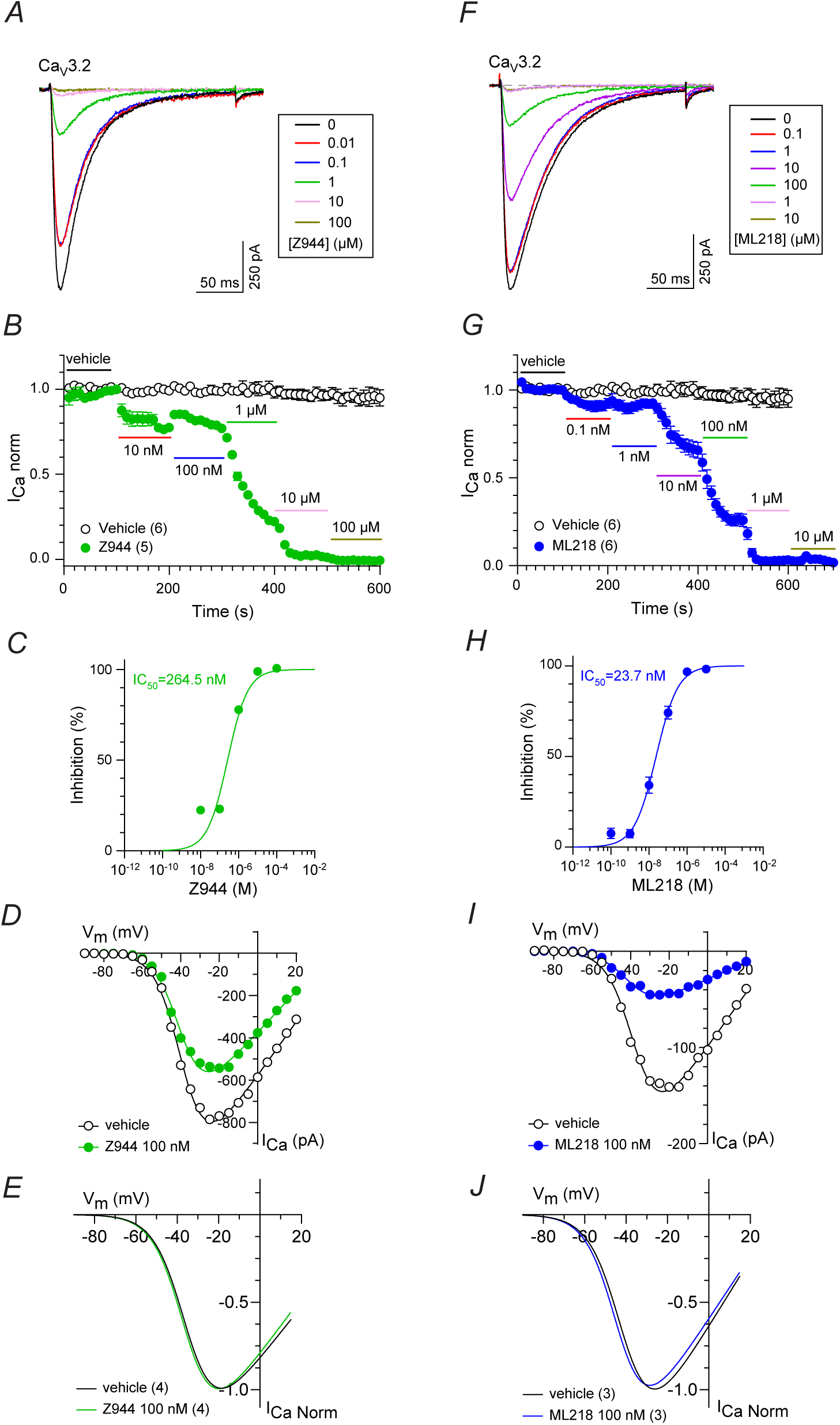
Inhibition of Ca_V_3.2 by Z944 and ML218. **(A)** Representative traces for Ca_V_3.2 *I_Ca_* elicited by 200-ms test pulses from a holding voltage of -90 mV to -20 mV before and after exposure to Z944. **(B)** Time course of *I_Ca_* treated with vehicle and increasing concentrations of Z944. *I_Ca norm_* represents current amplitude normalized to that measured during vehicle application. Points represented mean ± SEM. **(C)** For data in *B*, the inhibition (%) of *I_Ca_* was plotted as a function of Z944 concentration and fit by non-linear regression. **(D)** Representative current-voltage (I-V) relationship for *I_Ca_* before and after exposure to Z944. *I_Ca_* was evoked by 200-ms test pulses from -90 mV. **(E)** Boltzmann fits of I-V relationships of averaged data obtained as in *D,* were normalized to the peak *I_Ca_*. **(F-J)** Same as **A-E** but with ML218. Parentheses indicate numbers of cells.

**Figure 2.**
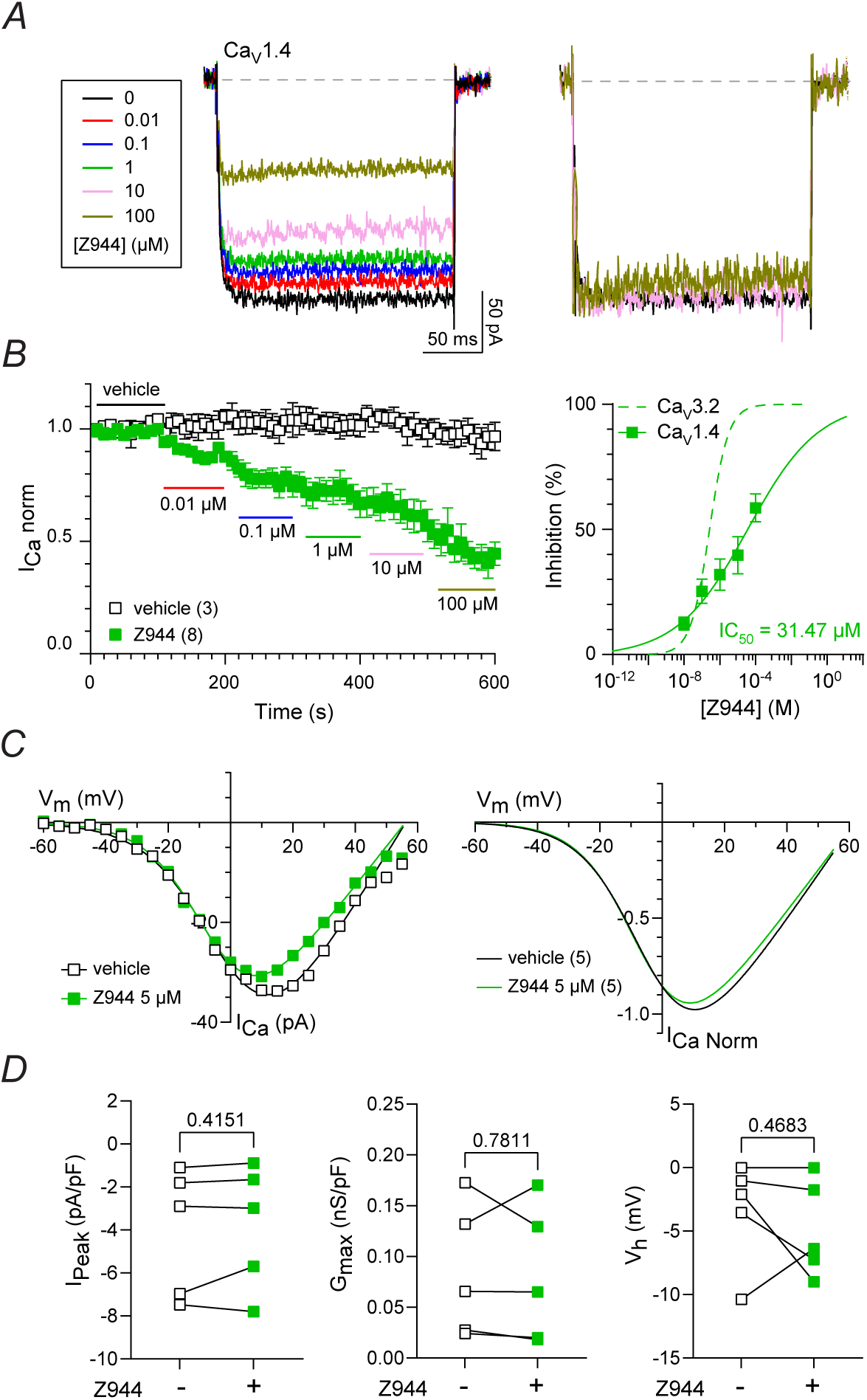
Inhibition of Ca_V_1.4 by Z944. **(A)** *Left*, representative traces for Ca_V_1.4 *I_Ca_* elicited by 200-ms test pulses from a holding voltage of -90 mV to +10 mV before and after exposure to Z944. *Right*, *I_Ca_* traces (normalized to vehicle-treated) show little to no inactivation with Z944. **(B)** *Left*, time course of *I_Ca_* treated with vehicle and increasing concentrations of Z944. *I_Ca norm_* represents current amplitude normalized to that measured during vehicle application. Points represent mean ± SEM. *Right*, for data in *B*, inhibition (%, normalized to *I_Ca_* during vehicle application) was plotted as a function of Z944 concentration and fit by non-linear regression. Dashed line represents curve fit of data for Ca_v_3.2 (Fig.1B). **(C)** *Left*, Representative current-voltage (I-V) relationship for *I_Ca_* before and after exposure to Z944. *I_Ca_* was evoked by 200-ms test pulses from -90 mV. *Right*, Boltzmann fits of I-V relationships of averaged data were normalized to the peak *I_Ca_*. **(D)** Peak current density (*I_Peak_*), *G_max_*, and *V_h_* before and after exposure to Z944 (5 µM). p-values were determined by paired t-tests. In *B,C*, parentheses indicate numbers of cells.

### Data presentation and statistical analysis

All data were compiled and analyzed statistically using GraphPad Prism software. Averaged data are presented as the mean ± SEM. Normality was assessed with the Shapiro–Wilk test. Parametric data were analyzed using the student’s t-test, while nonparametric data were analyzed using the Mann–Whitney or Wilcoxon tests. Normalized current-voltage relationship (I-V) data were fit to the Boltzmann equation (Eq. 1), where *G_max_* is the maximal conductance*, V_m_* is the test voltage*, V_rev_* is the apparent reversal potential*, V_h_* is the half maximal activation voltage, and *k* is the slope factor.

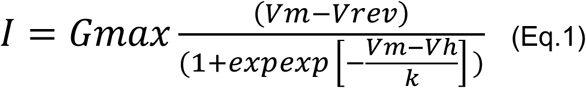

Dose–response curves were fitted using the [Inhibitor] vs. response least-square nonlinear regression equation (Eq.2). Where *X* is the concentration of the antagonist*, Y* is % inhibition*, IC_50_* is the half-maximal inhibitory concentration. Bottom and Top values were constrained to 0 and 100.

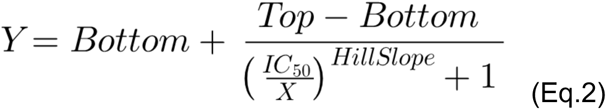

Comparisons between dose-response curve fits were made using the extra sum-of-squares F test.

### Computational modeling of human Ca_V_1.4 channel and ligand docking

The structural model of the human Ca_V_1.4 channel was generated using AlphaFold2 (AF2) (Jumper et al., 2021). Low-confidence unstructured regions of human Ca_V_1.4 predicted by AF2, including the N- and C-termini, were trimmed before docking preparation. To enable sampling of different conformational contexts around potential ligand-binding pockets of Ca_v_1.4, multiple models were generated using the following cryo-EM structures as templates: Ca_v_3.1 in complex with Z944 (PDB: 6kzp)(Zhao et al., 2019b), Ca_v_3.2 in complex with ML218 (PDB: 9ayk)(Huang et al., 2024), Ca_v_1.1 in complex with nifedipine (PDB: 6jp5)(Zhao et al., 2019a), and Ca_v_1.2 (PDB: 8eog)(Chen et al., 2023). All Ca_v_1.4 models differed in binding pocket geometry depending on the structural template used. Each Ca_v_1.4 model was then used for docking ML218 and Z944 into their respective binding contexts: the Ca_v_3-inspired fenestration, the canonical DHP site, and cavity adjacent to the DHP binding pocket identified by molecular surface analysis. Ligand docking was performed using both RosettaLigand (Meiler and Baker, 2006) and GALigandDock (Park et al., 2021) methods.

Supplementary Figures 1-3 show chemical structures and data supporting electrophysiological and modeling experiments and are included in Supplemental Material.

## RESULTS

### ML218 is more potent than Z944 in inhibiting Ca_v_3.2

The inhibitory actions of Z944 and ML218 on Ca_v_3.2 have been documented previously, but under varying recording conditions and in different cell lines (Xiang et al., 2011; Tringham et al., 2012; Huang et al., 2024). To rigorously compare the effects of these drugs on Ca_v_3.2 and Ca_v_1.4, we first compared their dose-response properties in HEK293T cells transfected with the Ca_v_ variants under identical experimental conditions. As anticipated, Z944 inhibited Ca_V_3.2 Ca^2+^ currents (*I_Ca_*; IC_50_∼ 265 nM, Fig.1A-C, Table 2). At 100 nM, Z944 caused ∼40% inhibition of the peak *I_Ca_* amplitude (Fig.1D; Supp Fig.2A), a significant decrease in the maximal conductance (*G_max_*), and hyperpolarizing shift in the reversal potential (*V_rev_*; Fig.1D,E, Table 3). Compared to Z944, ML218 exhibited 10-fold higher potency (IC_50_∼ 24 nM, Fig.1F-H, Table 2), causing ∼64% inhibition of the peak *I_Ca_* amplitude at 100 nM and slight reductions in *G_max_* and *V_rev_* that did not reach statistical significance (Fig. 1I,J, Table 3; Supp Fig. 2B). Our results reveal ML218 as a more potent antagonist of Ca_v_3.2 than Z944.

**Table 2.** Dose-response properties of Z944 and ML218.

| **Z944** | **IC<sub>50</sub> (μM)** | **Hill Slope** | **F (df)** | **p value** |
| --- | --- | --- | --- | --- |
| Ca<sub>v</sub>3.2 | 0.26 ± 0.04 | 0.82 ± 0.10 | F(2,61) = 60.47 | p < 0.001 |
| Ca<sub>v</sub>1.4 | 31.5 ± 21 | 0.22 ± 0.04 | -- | -- |
| **ML218** | **IC<sub>50</sub> (μM)** | **Hill Slope** | **F (df)** | **p value** |
| Ca<sub>v</sub>3.2 | 0.0237 ± 0.0025 | 0.75 ± 0.06 | F(2,62) = 144.7 | p < 0.001 |
| Ca<sub>v</sub>1.4 | 2.1 ± 0.51 | 0.46 ± 0.05 | -- | -- |
| Ca<sub>v</sub>1.4 T1007Y | 20 ± 4.9 | 0.57 ± 0.08 | F(2,51) = 21.74 | p < 0.001 |
| Ca<sub>v</sub>1.4 M1004A | 310 ± 190 | 0.24 ± 0.03 | F(2, 66) = 65.95 | p < 0.001 |
| Ca<sub>v</sub>1.4 M1129A | 0.45 ± 0.16 | 0.34 ± 0.05 | F(2, 45) = 7.61 | p < 0.002 |
| Ca<sub>v</sub>1.4 M1426A | 15 ± 2.9 | 0.72 ± 0.10 | F(2, 50) = 21.94 | p < 0.001 |
| Ca<sub>v</sub>1.2 | 37 ± 6.5 | 0.80 ± 0.11 | F(2,56) = 49.95 | p < 0.001 |
| Ca<sub>v</sub>1.2 V1063M | 120 ± 60 | 0.72 ± 0.29 | F(2, 56) = 3.96 | p = 0.025* |

**Table 3.**
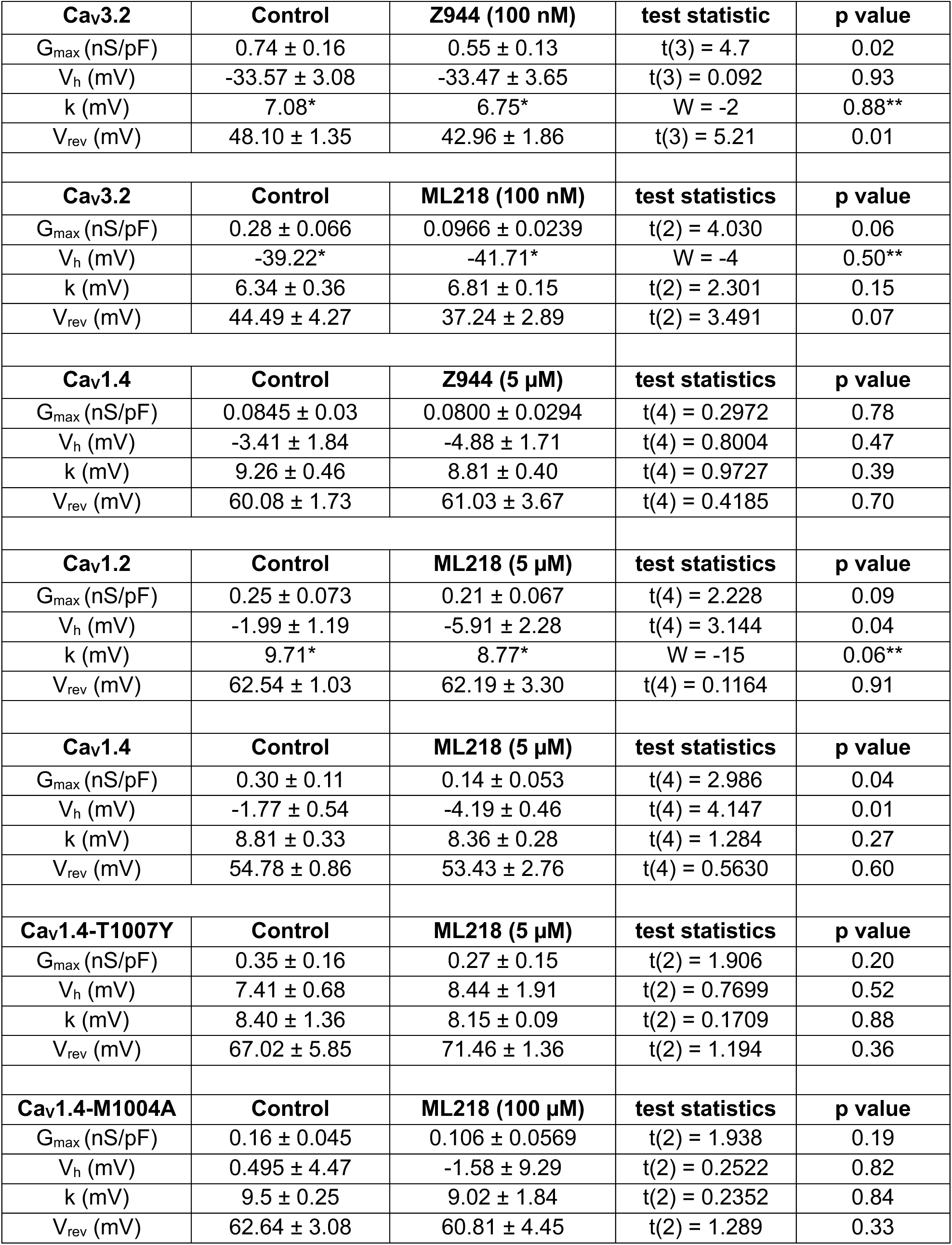
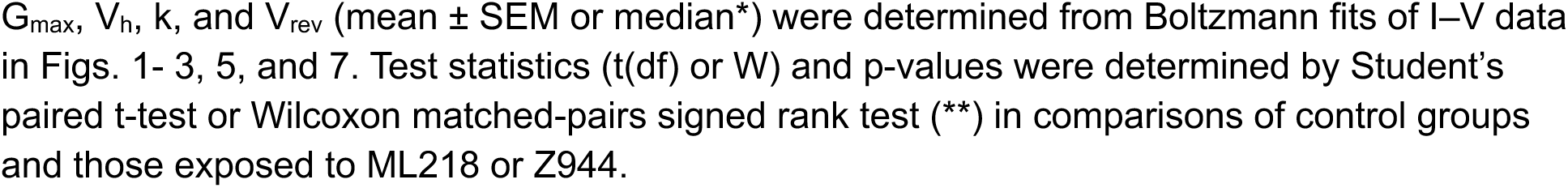
Effect of Z944 and ML218 on I-V parameters.

| **Ca<sub>v</sub>3.2** | **Control** | **Z944 (100 nM)** | **test statistic** | **p value** |
| --- | --- | --- | --- | --- |
| G<sub>max</sub> (nS/pF) | 0.74 ± 0.16 | 0.55 ± 0.13 | t(3) = 4.7 | 0.02 |
| V<sub>h</sub> (mV) | -33.57 ± 3.08 | -33.47 ± 3.65 | t(3) = 0.092 | 0.93 |
| k (mV) | 7.08* | 6.75* | W = -2 | 0.88** |
| V<sub>rev</sub> (mV) | 48.10 ± 1.35 | 42.96 ± 1.86 | t(3) = 5.21 | 0.01 |
| **Ca<sub>v</sub>3.2** | **Control** | **ML218 (100 nM)** | **test statistics** | **p value** |
| G<sub>max</sub> (nS/pF) | 0.28 ± 0.066 | 0.0966 ± 0.0239 | t(2) = 4.030 | 0.06 |
| V<sub>h</sub> (mV) | -39.22* | -41.71* | W = -4 | 0.50** |
| k (mV) | 6.34 ± 0.36 | 6.81 ± 0.15 | t(2) = 2.301 | 0.15 |
| V<sub>rev</sub> (mV) | 44.49 ± 4.27 | 37.24 ± 2.89 | t(2) = 3.491 | 0.07 |
| **Ca<sub>v</sub>1.4** | **Control** | **Z944 (5 μM)** | **test statistics** | **p value** |
| G<sub>max</sub> (nS/pF) | 0.0845 ± 0.03 | 0.0800 ± 0.0294 | t(4) = 0.2972 | 0.78 |
| V<sub>h</sub> (mV) | -3.41 ± 1.84 | -4.88 ± 1.71 | t(4) = 0.8004 | 0.47 |
| k (mV) | 9.26 ± 0.46 | 8.81 ± 0.40 | t(4) = 0.9727 | 0.39 |
| V<sub>rev</sub> (mV) | 60.08 ± 1.73 | 61.03 ± 3.67 | t(4) = 0.4185 | 0.70 |
| **Ca<sub>v</sub>1.2** | **Control** | **ML218 (5 μM)** | **test statistics** | **p value** |
| G<sub>max</sub> (nS/pF) | 0.25 ± 0.073 | 0.21 ± 0.067 | t(4) = 2.228 | 0.09 |
| V<sub>h</sub> (mV) | -1.99 ± 1.19 | -5.91 ± 2.28 | t(4) = 3.144 | 0.04 |
| k (mV) | 9.71* | 8.77* | W = -15 | 0.06** |
| V<sub>rev</sub> (mV) | 62.54 ± 1.03 | 62.19 ± 3.30 | t(4) = 0.1164 | 0.91 |
| **Ca<sub>v</sub>1.4** | **Control** | **ML218 (5 μM)** | **test statistics** | **p value** |
| G<sub>max</sub> (nS/pF) | 0.30 ± 0.11 | 0.14 ± 0.053 | t(4) = 2.986 | 0.04 |
| V<sub>h</sub> (mV) | -1.77 ± 0.54 | -4.19 ± 0.46 | t(4) = 4.147 | 0.01 |
| k (mV) | 8.81 ± 0.33 | 8.36 ± 0.28 | t(4) = 1.284 | 0.27 |
| V<sub>rev</sub> (mV) | 54.78 ± 0.86 | 53.43 ± 2.76 | t(4) = 0.5630 | 0.60 |
| **Ca<sub>v</sub>1.4-T1007Y** | **Control** | **ML218 (5 μM)** | **test statistics** | **p value** |
| G<sub>max</sub> (nS/pF) | 0.35 ± 0.16 | 0.27 ± 0.15 | t(2) = 1.906 | 0.20 |
| V<sub>h</sub> (mV) | 7.41 ± 0.68 | 8.44 ± 1.91 | t(2) = 0.7699 | 0.52 |
| k (mV) | 8.40 ± 1.36 | 8.15 ± 0.09 | t(2) = 0.1709 | 0.88 |
| V<sub>rev</sub> (mV) | 67.02 ± 5.85 | 71.46 ± 1.36 | t(2) = 1.194 | 0.36 |
| **Ca<sub>v</sub>1.4-M1004A** | **Control** | **ML218 (100 μM)** | **test statistics** | **p value** |
| G<sub>max</sub> (nS/pF) | 0.16 ± 0.045 | 0.106 ± 0.0569 | t(2) = 1.938 | 0.19 |
| V<sub>h</sub> (mV) | 0.495 ± 4.47 | -1.58 ± 9.29 | t(2) = 0.2522 | 0.82 |
| k (mV) | 9.5 ± 0.25 | 9.02 ± 1.84 | t(2) = 0.2352 | 0.84 |
| V<sub>rev</sub> (mV) | 62.64 ± 3.08 | 60.81 ± 4.45 | t(2) = 1.289 | 0.33 |

### ML218 is more potent than Z944 in inhibiting Ca_V_1.4 channels

To follow-up on prior reports of Z944 and ML218 in modulating Ca_v_1.4 channels in mouse cones (Davison et al., 2022; Maddox et al., 2024), we analyzed their effects in HEK293T cells transfected with Ca_V_1.4 along with the auxiliary β_2×13_ and α_2_δ-4 subunits which co-assemble with Ca_v_1.4 in the retina (Lee et al., 2015). Consistent with its effects on Ca_v_1.2 (Tringham et al., 2012), Z944 inhibited Ca_v_1.4 with an IC_50_ (∼31 µM) that was ∼100-fold higher than that for Ca_v_3.2 (Fig. 2A,B, Table 2). At relatively high concentrations (10 and 100 µM), Z944 had little impact on the slow inactivation that is characteristic of Ca_v_1.4 (Fig.2A). At 5 µM, Z944 had no significant effect on *I_Ca_* amplitude or other parameters of the I-V relationship (Fig. 2C,D; Table 3).

In contrast to the modest effects of Z944, ML218 robustly inhibited Ca_V_1.4 currents at low micromolar doses (IC_50_ ∼ 2 µM; Table 1, Fig. 3A,B). At 5 µM, ML218 caused greater than 50% inhibition of *I_Ca_*, decreased *G_max_*, and hyperpolarized the *V_h_* (Fig.3C,D; Table 3; Supp Fig. 2D). The inhibitory and stimulatory effects of ML218 are reminiscent of the dual agonist/antagonist properties of racemic mixtures of DHP derivatives (Schramm et al., 1983b; Franckowiak et al., 1985). Moreover, ML218 (10 and 100 μM) increased inactivation of Ca_v_1.4 *I_Ca_* (Fig.3A), like some DHP antagonists (Berjukow et al., 2000). Contrary to the enhanced block of Ca_v_1 channels by some DHPs at depolarized voltages (Welling et al., 1997; Koschak et al., 2001; Koschak et al., 2003), Ca_v_1.4 inhibition by ML218 was similar whether the holding voltage was set to -90 mV or -50 mV (Fig.4).

**Figure 3.**
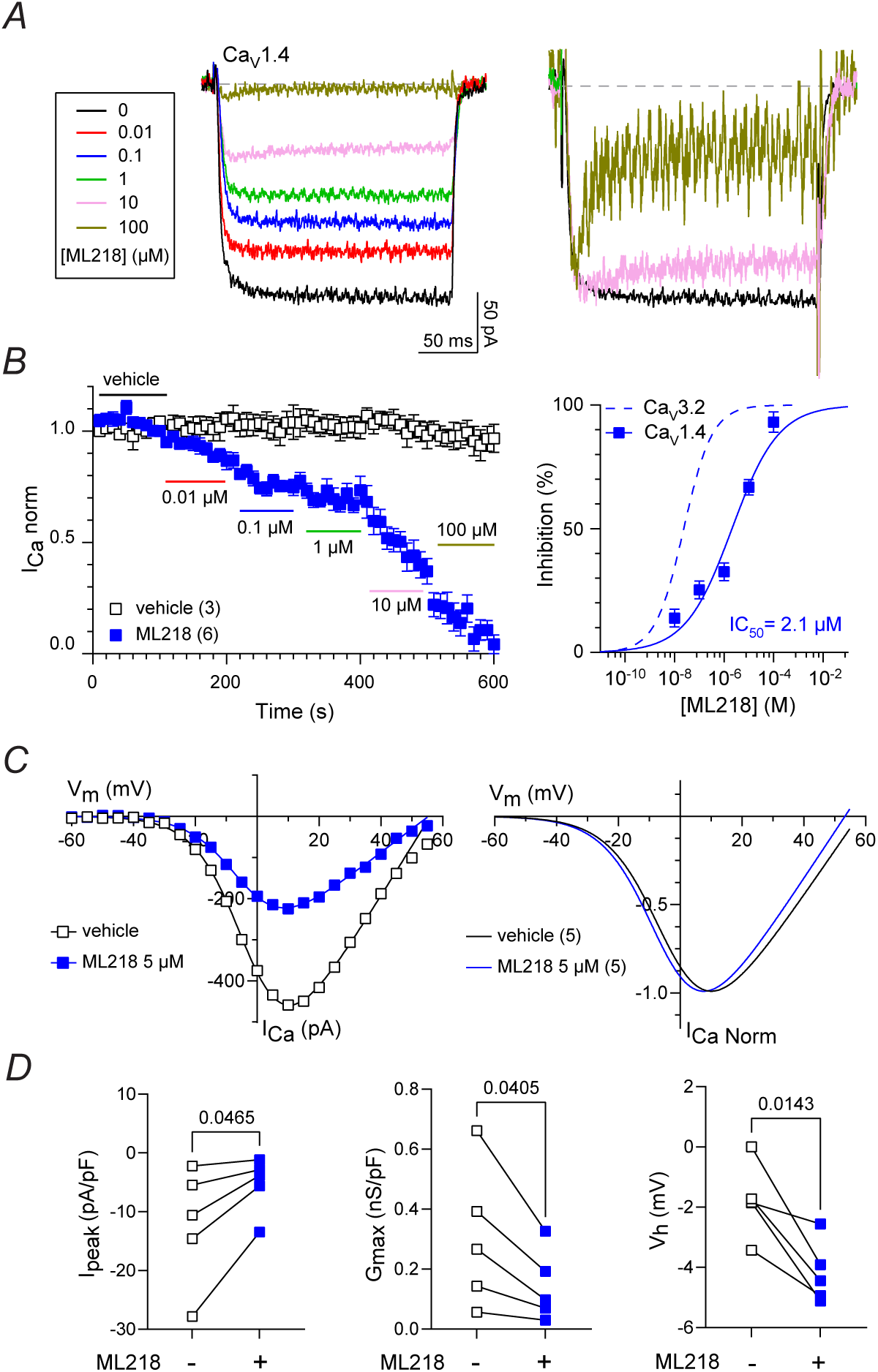
Inhibition of Ca_V_1.4 by ML218. **(A)** *Left*, representative traces for Ca_V_1.4 *I_Ca_* elicited by a 200-ms test pulse from -90 mV to +10 mV before and after exposure to ML218. *Right*, *I_Ca_* traces (normalized to vehicle-treated) show inactivation with ML218 at 10 and 100 μM. **(B)** *Left*, time course of *I_Ca_* treated with vehicle (control) or the indicated concentrations of ML218. *I_Ca norm_* represents current amplitude normalized to that measured during vehicle application. Points represent mean ± SEM. *Right*, Dose-response plot showing the inhibition (%) of *I_Ca_* as a function of ML218 concentration fit by non-linear regression. Dashed line represents curve fit of data for Ca_v_3.2 (Fig.1H). **(C)** *Left*, representative I-V plot for *I_Ca_* before and after exposure to ML218. *I_Ca_* was evoked by 200-ms test pulses from -90 mV. Smooth lines represent Boltzmann fits. *Right,* Boltzmann fits of I-V data normalized to the peak *I_Ca_* **(D)** Peak current density (I_Peak_), normalized G_max_, and V_h_ before and after exposure to 5 µM ML218. p-values were determined by paired t-tests. In *B,C*, parentheses indicate numbers of cells.

**Figure 4.**
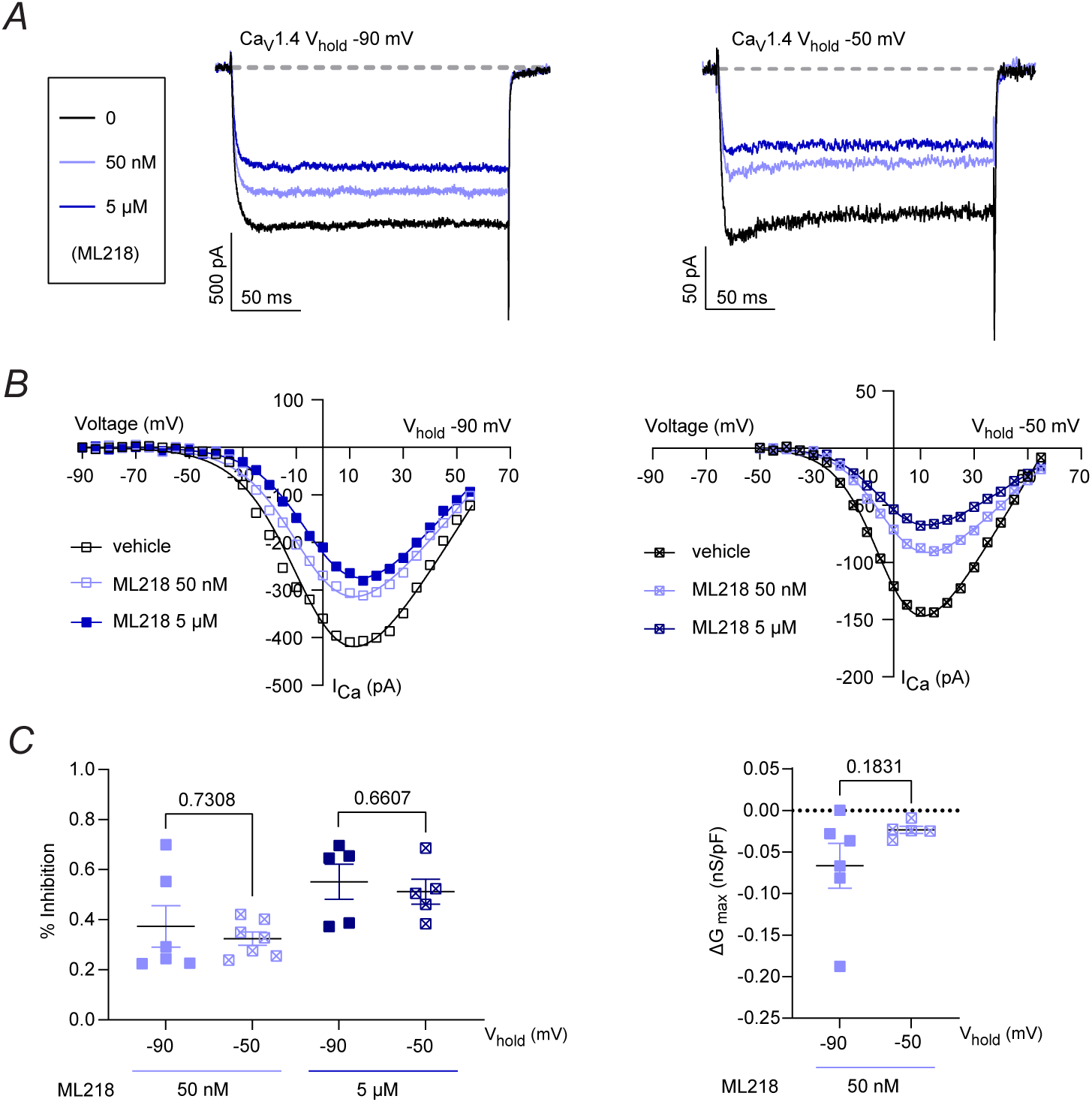
Inhibition of Ca_V_1.4 by ML218 is voltage-independent. **(A)** *R*epresentative traces for Ca_V_1.2 *I_Ca_* elicited by a 200-ms test pulse from holding voltage (V_hold_) of -90 mV or -50 mV to +10 mV before and after exposure to ML218. **(B)** Representative I-V plots for *I_Ca_* evoked by 50-ms test pulses from holding voltage (V_hold_) of -90 mV or -50 mV before and after exposure to ML218. Smooth lines represent Boltzmann fits. **(C)** Inhibition (%) and difference in *G_max_* (ΔG_max_) caused by ML218 (5 µM). p-values were determined by unpaired t-tests.

We next tested whether ML218 could modulate another DHP-sensitive Ca_v_1 channel, Ca_v_1.2. In contrast to its effects on Ca_v_1.4, ML218 inhibited Ca_v_1.2 only at high micromolar concentrations (IC_50_∼37 µM; Fig.5A,B, Table 2). While it strongly enhanced inactivation at 100 µM (Fig.5A), ML218 at 5 µM did not significantly inhibit Ca_V_1.2 *I_Ca_* or *G_max_* (Fig.5C,D) but hyperpolarized *V_h_*, as observed for Ca_V_1.4 (Fig. 5C,D; Table 3). Our results indicate that Z944 and ML218 can modulate Ca_v_1.4, and that ML218 has effects that are similar to DHPs with higher potency for Ca_v_1.4 than for Ca_v_1.2.

**Figure 5.**
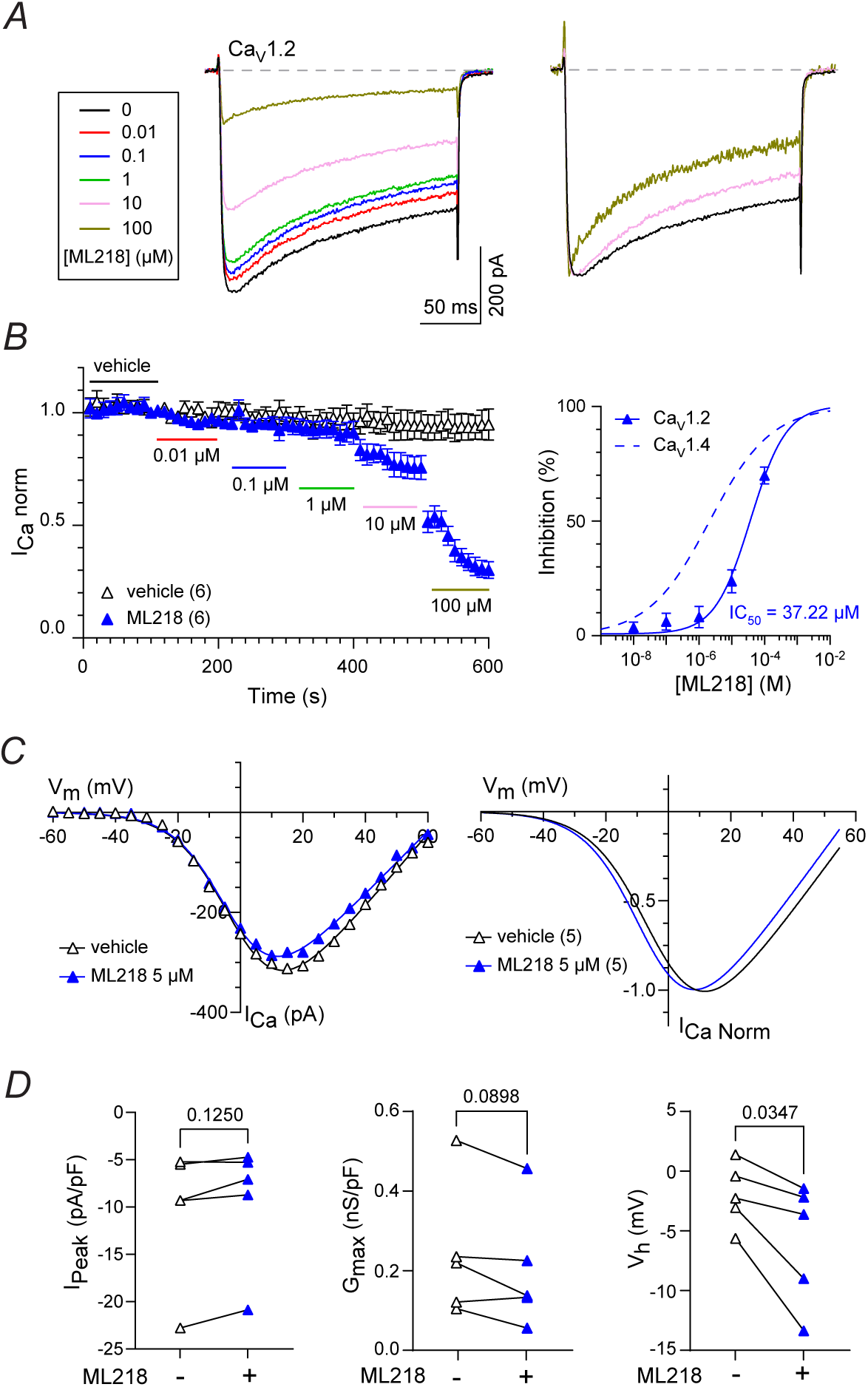
Inhibition of Ca_V_1.2 by ML218. **(A)** *Left*, representative traces for Ca_V_1.2 *I_Ca_* before and after exposure to ML218. *I_Ca_* was elicited by a 200-ms test pulse from -90 mV to 10 mV. *Right*, *I_Ca_* traces (normalized to vehicle-treated) show inactivation with ML218 at 10 and 100 μM. **(B)** *Left*, time course of *I_Ca_* treated with vehicle (control) or the indicated concentrations of ML218. *I_Ca norm_* represents current amplitude normalized to that measured during vehicle application. Points represent mean ± SEM. *Right*, Dose-response plot showing the inhibition (%) of *I_Ca_* as a function of ML218 concentration fit by non-linear regression. Dashed line represents curve fit of data for Ca_v_1.4 (Fig.3B). **(C)** *Left*, representative I-V plot for *I_Ca_* before and after exposure to ML218. *I_Ca_* was evoked by 200-ms test pulses from -90 mV. Smooth lines represent Boltzmann fits. *Right,* Boltzmann fits of I-V data normalized to the peak *I_Ca_* **(D)** Peak current density (I_Peak_), normalized G_max_, and V_h_ before and after exposure to 5 µM ML218. p-values were determined by paired t-tests. In *B,C*, parentheses indicate numbers of cells.

### Role of the DHP binding site in the effects of ML218 on Ca_v_1 channels

The pore-forming α_1_ subunit of Ca_v_1.4 consists of four homologous domains, each with six transmembrane helices (S1-S6). An extracellular membrane-re-entrant P-loop (P1, P2) forms the selectivity-filter between helices S5 and S6. S1–S4 form a voltage-sensing domain, while S5, S6, and the P-loop form to the pore domain (Fig.6A). Structural, functional, and biochemical studies indicate that DHPs bind to a fenestration formed by the pore-forming segments in domains III and IV (IIIS5, IIIP1, IIIS6, and IVS6 (Striessnig et al., 1991; Peterson et al., 1996; Schuster et al., 1996; Peterson et al., 1997; Yamaguchi et al., 2000; Zhao et al., 2019a)). A threonine residue in IIIS5, which is conserved in all Ca_v_1 subtypes (T1007 in Ca_v_1.4, Fig.6A), forms a hydrogen bond with the C3 ester of DHPs (Zhao et al., 2019a). The sensitivity of Ca_v_1.2 to DHPs is blunted upon mutating this threonine to the tyrosine residue present in Ca_v_2 channels at this position (Mitterdorfer et al., 1996; Sinnegger-Brauns et al., 2004). To test whether the actions of ML218 on Ca_v_1.4 required the DHP binding site, we mutated T1007 to tyrosine (Ca_v_1.4-T1007Y).

**Figure 6.**
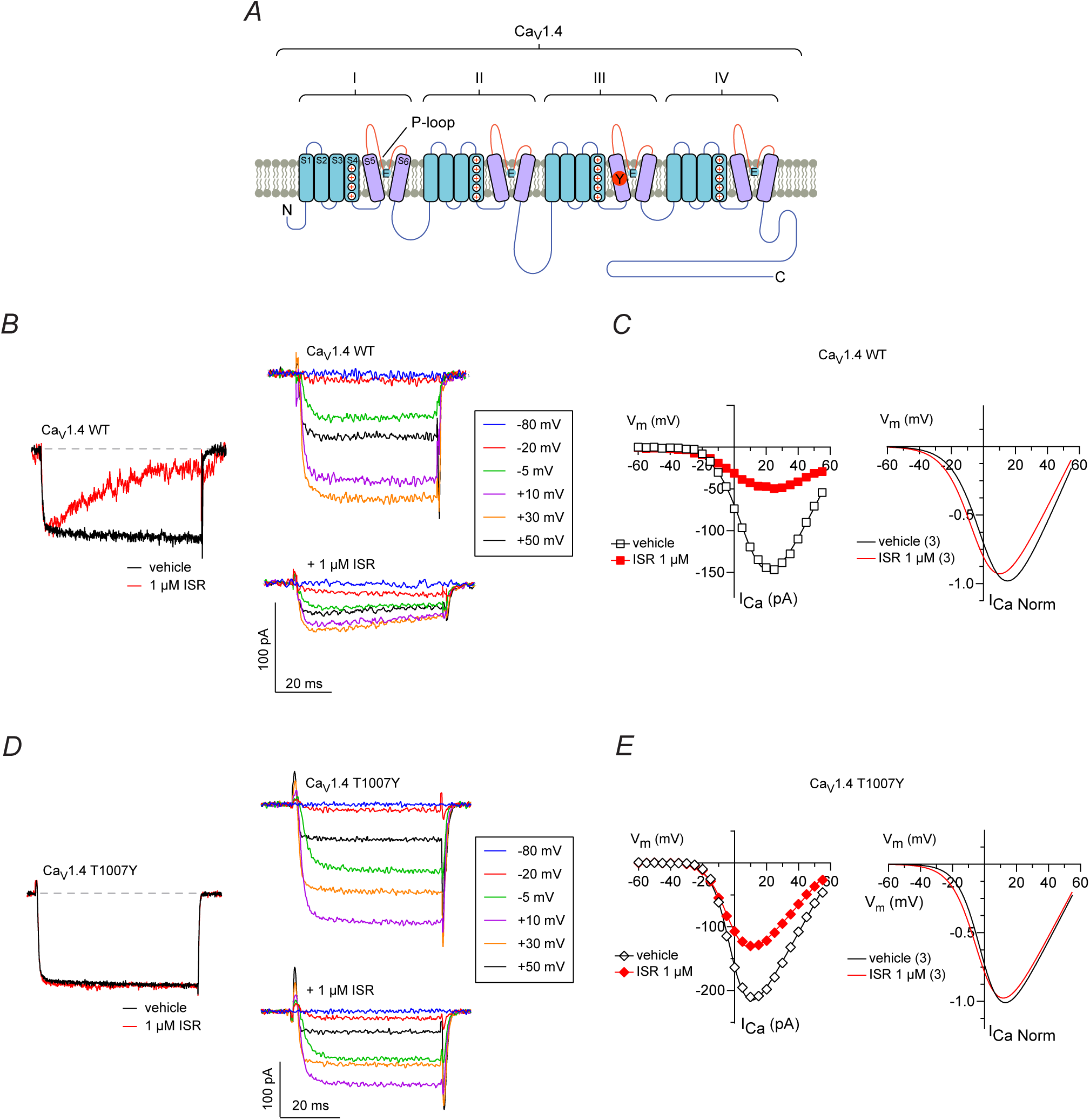
Effect of mutation of T1007Y on Ca_v_1.4 inhibition by isradipine. **(A)** Schematic showing Ca_V_1.4 with four repeats (I-IV) each with 6 transmembrane helices (S1-S6) and pore (P) loop. T1007Y mutation in IIIS5 is indicated (red circle). **(B)** *Left*, Ca_V_1.4 WT *I_Ca_* traces (normalized to vehicle-treated) show inactivation with ISR. *I_Ca_* was elicited by a 200-ms test pulse from -90 mV to 10 mV. *Right*, representative *I_Ca_* traces of Ca_V_1.4 WT before and after exposure to isradipine (ISR). *I_Ca_* was elicited by a 50-ms test pulse from -90 mV to indicated voltages. **(C)** *Left*, representative I-V plot for *I_Ca_* before and after exposure to ISR. *I_Ca_* was evoked by 50-ms test pulses from a holding potential of -90 mV. Smooth lines represent Boltzmann fits. *Right,* Boltzmann fits of I-V data normalized to the peak *I_Ca_* (n = 3). **(D,E)** Same as *B,C,* but for Ca_V_1.4 T1007Y.

We first confirmed that Ca_v_1.4-T1007Y exhibited a weakened sensitivity to the DHP antagonist, isradipine (ISR). For wild type (WT) Ca_v_1.4, ISR (1 μM) produced a ∼78% inhibition of *I_Ca_* amplitude and greatly enhanced inactivation (Fig.6B,C; Supp Fig. 2G). While having less of an impact in Ca_v_1.4 than Ca_v_1.2 (Mitterdorfer et al., 1996; Sinnegger-Brauns et al., 2004), the T1007Y mutation significantly reduced the response to ISR. ISR (1 μM) caused only ∼17 % inhibition of Ca_v_1.4-T1007Y *I_Ca_* and did not affect inactivation (Fig.6D,E; Supp Fig. 2H). Ca_v_1.4-T1007Y was also less sensitive to the effect of ML218. The IC_50_ for Ca_v_1.4-T1007Y (∼20 µM) was 10-fold higher than that for Ca_v_1.4-WT (Fig. 7A,B; Table 1). At 5 µM, ML218 produced only ∼33% inhibition of Ca_v_1.4-T1007Y *I_Ca_* compared to ∼50% for Ca_v_1.4 WT, with no effects on *G_max_* or *V_h_* (Fig. 7C,D; Table 3; Supp Fig. 2I). These results support a role for the DHP binding pocket in the modulatory actions of ML218 on Ca_v_1.4.

**Figure 7.**
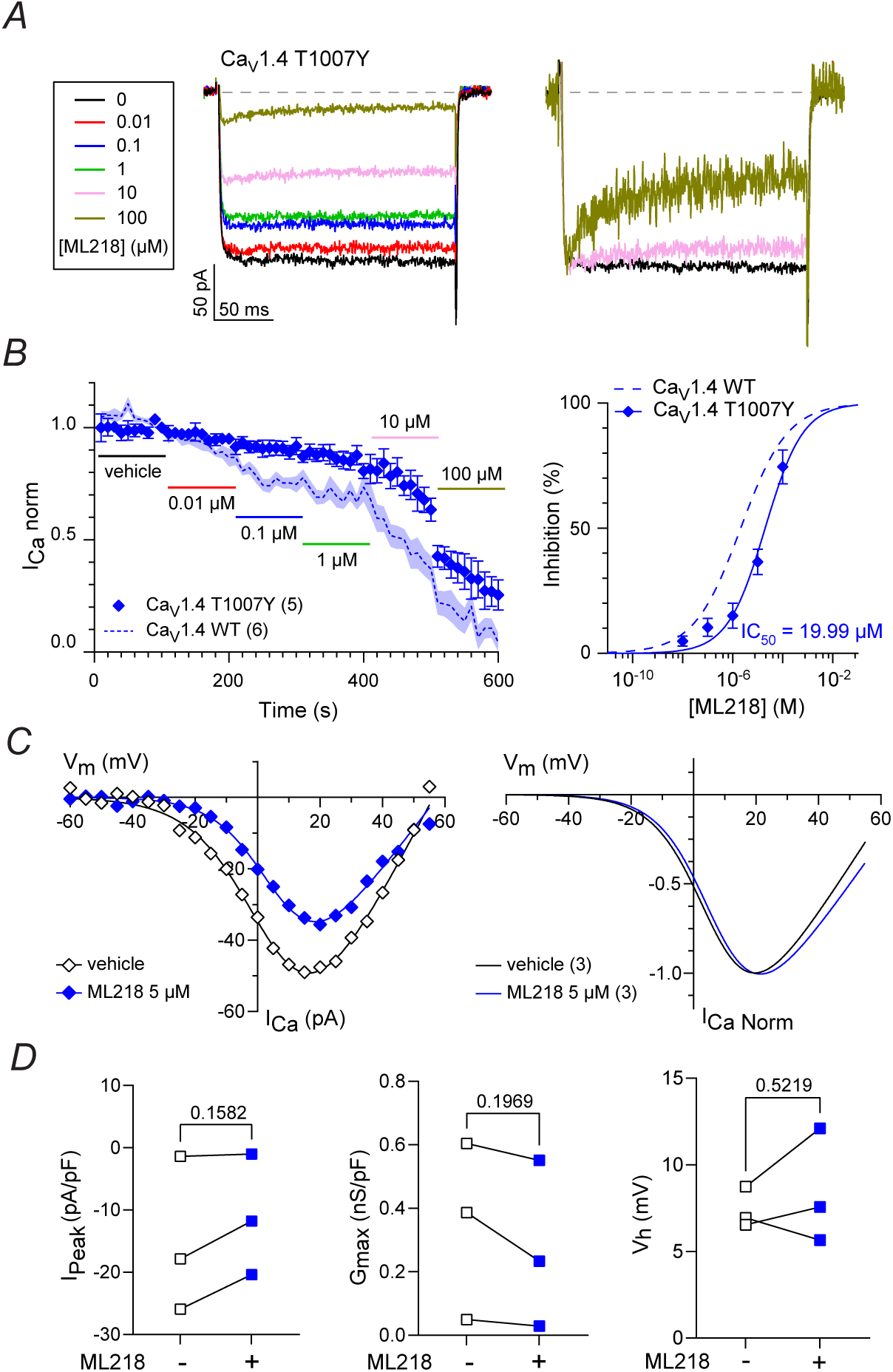
Effect of T1007Y mutation on Ca_v_1.4 inhibition by ML218. **(A)** *Left*, representative Ca_v_1.4 T1007Y *I_Ca_* traces before and after exposure to ML218. *I_Ca_* was elicited by a 200-ms test pulse from -90 mV to +10 mV. *Right*, *I_Ca_* traces (normalized to vehicle-treated) show inactivation with ML218 at 10 and 100 μM. **(B)** *Left*, time course of *I_Ca_* treated with vehicle (control) or the indicated concentrations of ML218. *I_Ca norm_* represents current amplitude normalized to that measured during vehicle application. Points represent mean ± SEM. *Right*, Dose-response plot showing the inhibition (%) of *I_Ca_* as a function of ML218 concentration fit by non-linear regression. Dashed line represents curve fit of data for Ca_v_1.4 WT (Fig.3B). **(C)** *Left*, representative I-V plot for *I_Ca_* before and after exposure to ML218. *I_Ca_* was evoked by 200-ms test pulses from -90 mV. Smooth lines represent Boltzmann fits. *Right,* Boltzmann fits of I-V data normalized to the peak *I_Ca_* **(D)** Peak current density (*I_Peak_*), *G_max_*, and *V_h_* before and after exposure to ML218 (5 µM). p-values were determined by paired t-tests. In *B,C*, parentheses indicate numbers of cells.

### Structural modeling of ML218 and Z944 interactions with Ca_v_1.4

Cryo-electron microscopy (cryo-EM) has revealed structural details of ML218 and Z944 (Supp. Fig.3) binding to Ca_v_3 channels, with both drugs blocking the pore through binding within the I-IV and II-III fenestration and central cavity (Zhao et al., 2019b; Huang et al., 2024). Using AlphaFold 2 (AF2) (Jumper et al., 2021)), we generated a model of the Ca_v_1.4 pore and aligned the structures of ML218 and Z944 bound to Ca_V_3.2 and Ca_V_3.1 (Zhao et al., 2019b; Huang et al., 2024), respectively (Fig.8A,B). Aligning cryo-EM structures of Ca_V_3.2 and Ca_V_3.1 with the Ca_V_1.4 model reveals that several residues that form the ML218 and Z944 binding pocket are not conserved (Supp.Fig.3A,B).

**Figure 8.**
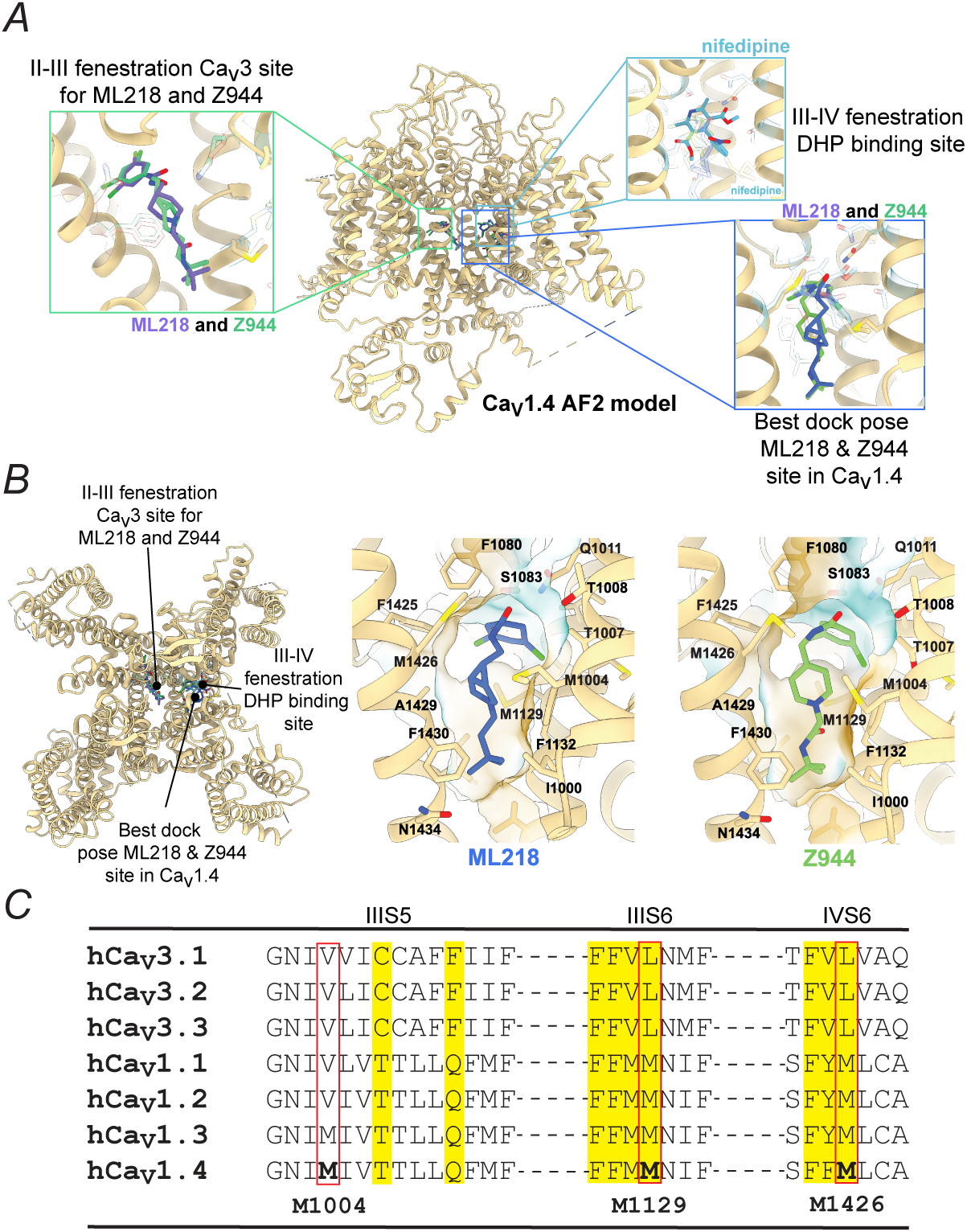
Modeling of ML218 and Z944 in Ca_V_1.4. **(A)** Structural overview of the Ca_V_1.4 α₁ subunit model generated by AlphaFold2, superimposed with experimental structures of nifedipine-bound rabbit Ca_V_1.1 (PDB:6jp5), ML218-bound Ca_V_3.2 (PDB: 9ayk), and Z944-bound Ca_V_3.1 (PDB: 6kzp). Boxed images show poses of ML218 and Z944 in the site corresponding to the Ca_v_3 binding pocket in the II-III fenestration, nifedipine binding in the canonical DHP site, and the best docking pose of ML218 and Z944 in the III-IV fenestration. **(B)** *Left panel,* top-down view of the Ca_V_1.4 model, showing the positions of the ligands in experimental structures (nifedipine, ML218, and Z944. *Middle and right panels*, best docking poses for ML218 and Z944 in the III-IV fenestration in Ca_V_1.4, highlighting key interacting residues and hydrophobic surfaces (green shading). **(C)** Alignment of residues in IIIS5, IIIS6, and IVS6 in Ca_v_3 and Ca_v_1 subtypes. Residues involved in DHP binding are highlighted. Numbers correspond to human Ca_V_1.4 sequence. ML218 and Z944 interacting methionine residues are in bold.

The docking of ML218 and Z944 in the Ca_V_1.4 II-III fenestration, using as templates the structures of Cav3.2 and Cav3.1, respectively, showed that neither ligand reproduced its Ca_v_3 binding mode in this region of Ca_V_1.4. The Rosetta ligand docking plots also did not display funnel-like behavior (Supp.Fig.3C,D), which is likely due to differences in binding pocket residues between Ca_v_1.4 and Ca_v_3 channels (Zhao et al., 2019b; Huang et al., 2024).

Considering our electrophysiological data, we next docked ML218 and Z944 into Ca_V_1.4 models generated with templates of Ca_V_1.1 and Ca_V_1.2, focusing on the DHP binding site within the III–IV fenestration (Fig.8A,B). Both ligands adopted favorable and overlapping poses in this region with their phenyl headgroups pointing towards the conserved Ca_V_1.4 residues T1007 and Q1011, which confer the specificity of DHP interactions with Ca_v_1 vs Ca_v_2 subtypes (Mitterdorfer et al., 1996; Hockerman et al., 1997; Sinnegger et al., 1997). The aliphatic tails of both ML218 and Z944 are positioned in the hydrophobic pocket of IIIS6 and IVS6 (Fig. 8B,C)(Zhao et al., 2019a; Wei et al., 2024).

For ML218, the proposed pose had a ligand interface score of –33.19 Rosetta Energy Units (REU) and displayed a clear funnel-like behavior (Supp.Fig.3F). For Z944, a low root mean square deviation (RMSD) relative to ML218 was observed, together with a favorable score of -34.32 REUs, suggesting that both ligands can similarly engage this novel binding pocket. Some alternate ML218 poses scored as low as –38.3 REU, but their Rosetta ligand docking plots were inconclusive, where poses that represented a rotated orientation of the molecule scored similar ligand interface scores (Supp.Fig.3E). These results suggest that the III-IV fenestration of Ca_v_1.4 could accommodate multiple binding modes for ML218.

### Methionine residues in the III-IV fenestration contribute to sensitivity of Ca_V_1.4 to ML218

The best docking poses for ML218 and Z944 involved a cluster of methionine residues (M1004, M1129, M1426; Fig.8). To test their functional relevance, we substituted them individually with alanine (M1004A, M1129A, M1426A). Like the T1007 mutation, the M-A mutations had nominal effects on the I-V parameters (Table 4). Whereas the M1004A and M1426A mutants were markedly less sensitive to ML218 than Ca_v_1.4 WT, the M1129A mutant was slightly more sensitive to ML218 (Fig.9, Table 2). Compared to Ca_v_1.4 WT, the inhibition by 10 μM ML218 of M1004A and M1426A mutants was ∼50% weaker, while that for M1129A was unchanged (Fig.9G). While M1129 and M1426 are conserved among all Ca_v_1 subtypes, M1004 is present in IIIS5 of Ca_v_1.4 and the closely related Ca_v_1.3 but corresponds to a valine in Ca_v_1.2 and Ca_v_1.1 (Fig.8C). If this conservative substitution contributed to the lower ML218 sensitivity of Ca_v_1.2 than of Ca_v_1.4, then mutating this site to methionine (V1063M) should increase the ML218 potency for Ca_v_1.2. However, V1063M actually increased the IC_50_ and did not increase inhibition of *I_Ca_* by 100 μM ML218 compared to the wild-type Ca_v_1.2 (Fig. 10C; Table 3). These results suggest that V1063 contributes to ML218 binding but cannot account for the lower sensitivity of Ca_v_1.2 to ML218 as compared to Ca_v_1.4.

**Figure 9.**
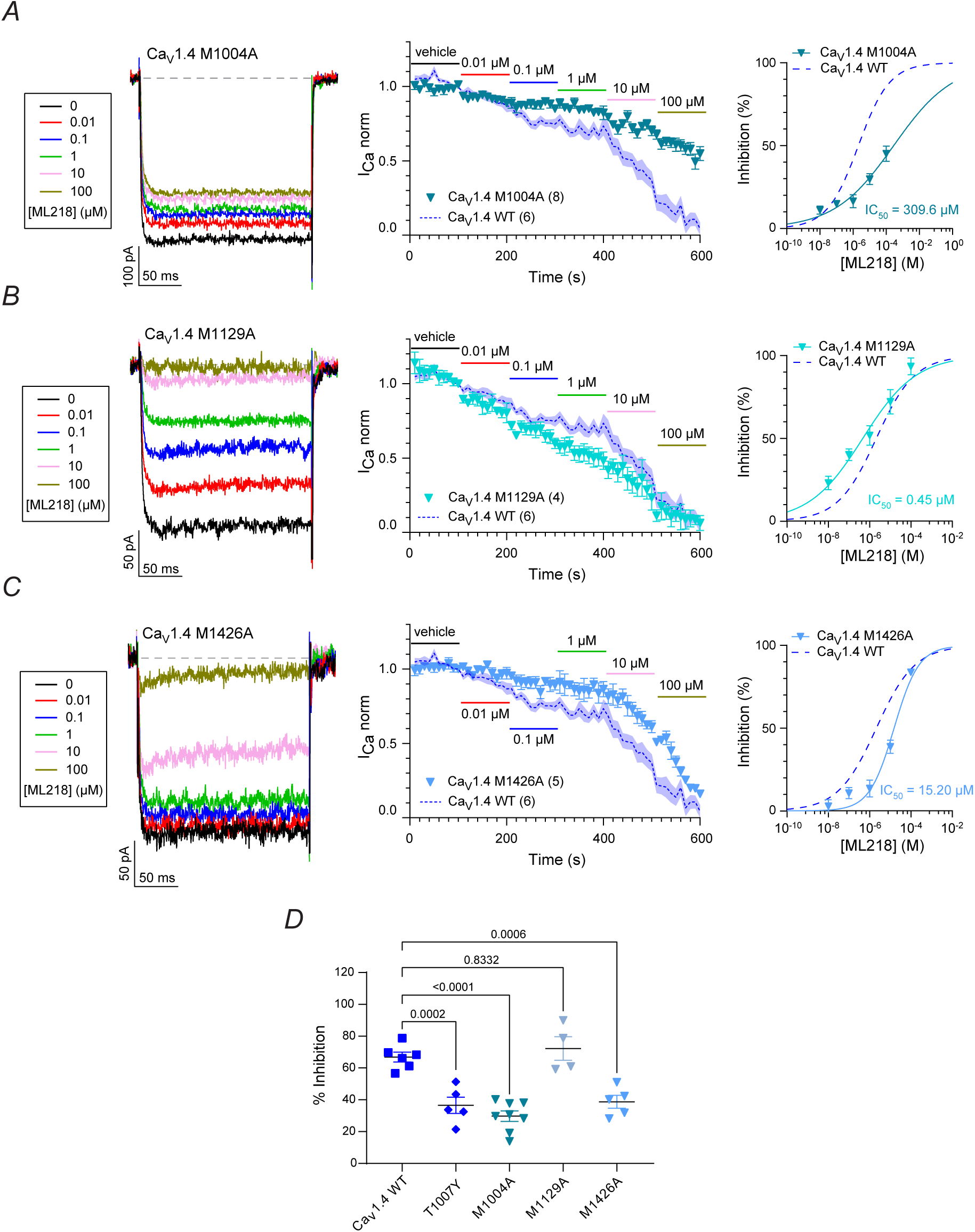
Effect of M1004A, M1129A, and M1426A mutations on Ca_v_1.4 inhibition by ML218. (A-C) *Left*, representative Ca_v_1.4 mutant *I_Ca_* traces before and after exposure to ML218. *I_Ca_* was elicited by a 200-ms test pulse from -90 mV to +10 mV. *Middle*, time course of *I_Ca_* treated with vehicle (control) or the indicated concentrations of ML218. *I_Ca norm_* represents current amplitude normalized to that measured during vehicle application. Points represent mean ± SEM. *Right*, Dose-response plot showing the inhibition (%) of *I_Ca_* as a function of ML218 concentration fit by non-linear regression. Dashed line represents curve fit of data for Ca_v_1.4 WT (Fig.3B). Parentheses indicate numbers of cells.. **(D)** % inhibition by ML218 (10 μM) of peak *I_Ca_* measured in middle panels of *A-C*. Bars represent mean ± SEM, p-value determined via Dunnett’s multiple comparisons test.

**Figure 10.**
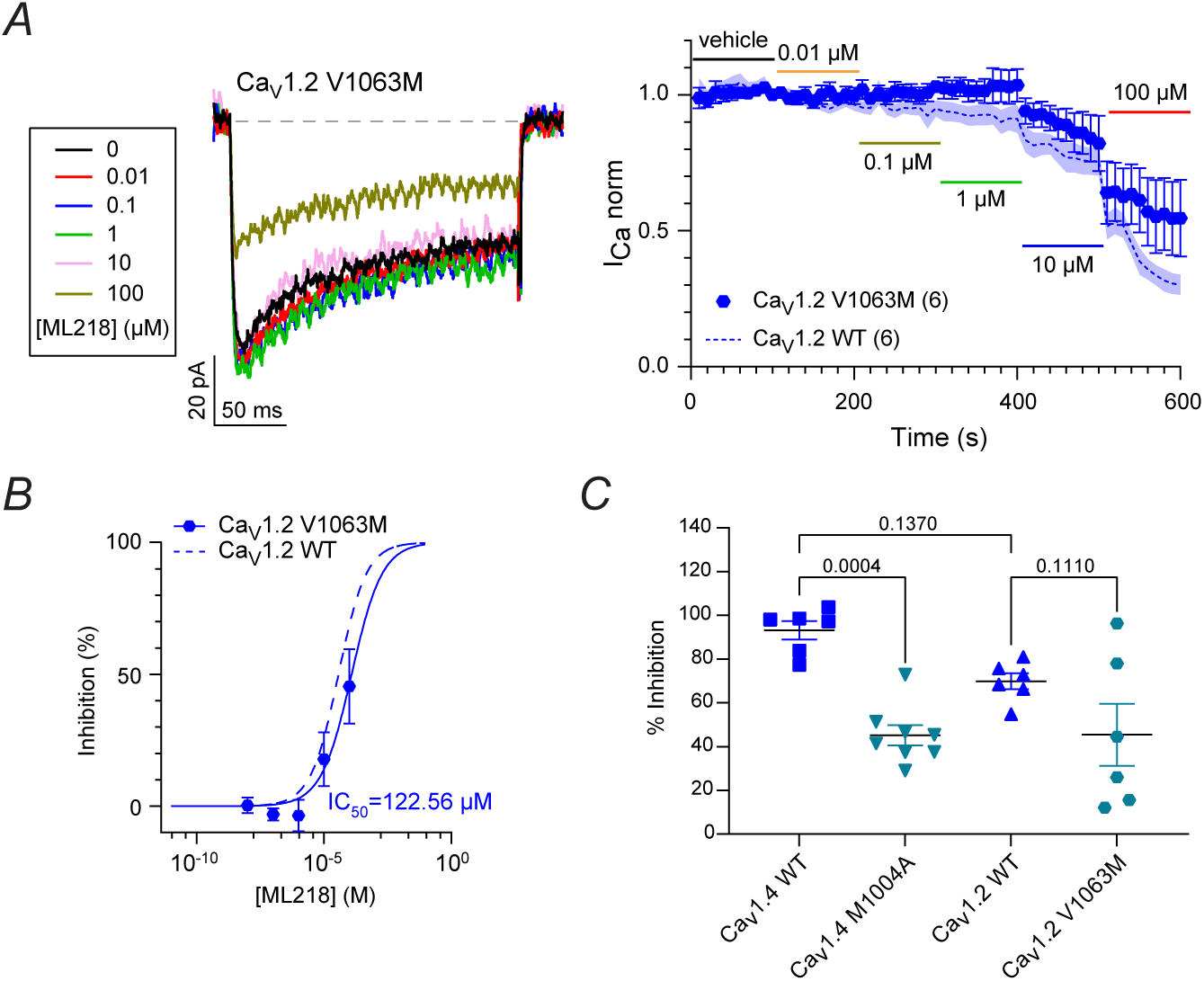
Effect of V1063M mutation on ML218 inhibition of Ca_v_1.2. *Left*, representative Ca_v_1.2 V1063M *I_Ca_* traces before and after exposure to ML218. *I_Ca_* was elicited by a 200-ms test pulse from -90 mV to +10 mV. *Right*, time course of *I_Ca_* treated with vehicle (control) or the indicated concentrations of ML218. *I_Ca norm_* represents current amplitude normalized to that measured during vehicle application. Points represent mean ± SEM. Parentheses indicate numbers of cells. **(B)** Dose-response plot showing the inhibition (%) of *I_Ca_* as a function of ML218 concentration fit by non-linear regression. Dashed line represents curve fit of data for Ca_v_1.2 WT (Fig.5B). **(C)** % inhibition by ML218 (100 μM) of peak *I_Ca_* measured in the time course panel of *A*. Bars represent mean ± SEM, p-value determined via Dunnett’s multiple comparisons test.

**Table 4.**
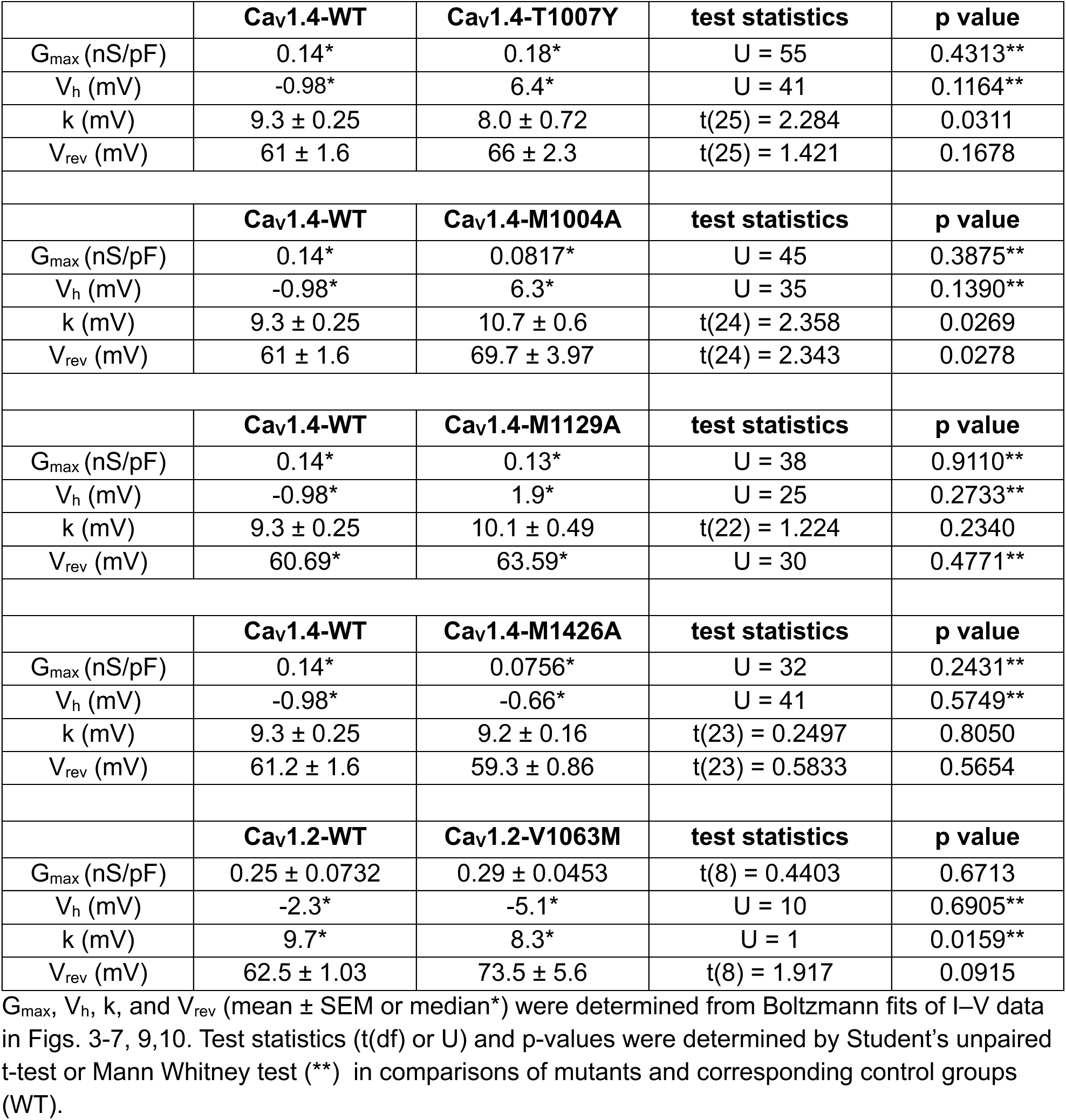
Effect of point mutations on I-V parameters.

## Discussion

Our study provides new insights into the molecular regulation of Ca_v_1 channels by pharmacological ligands. First, at concentrations typically used to experimentally block Ca_v_3 channels, Z944 and ML218 have additional modulatory actions on Ca_v_1.4. ML218 is more potent than Z944 as a Ca_v_1.4 antagonist and exhibits modest agonist activity. Second, our modeling and functional studies suggest that Z944 and ML218 exploit a binding site in domains III and IV of Ca_v_1.4 that includes residues of the DHP binding pocket. Third, a cluster of methionine residues within the III-IV fenestration contributes to the high potency of ML218 for Ca_v_1.4, but differences outside of this region are likely to account for the relatively weak antagonism of Ca_v_1.2 by ML218. We propose that ML218 and Z944 represent dual Ca_v_1/Ca_v_3 modulators which could be modified to achieve distinct selectivity profiles for different Ca_v_ subtypes.

### Binding modes of Ca_v_3/Ca_v_1 blockers

ML218 and Z944 have an elongated structure (Supp. Fig.1A), with a phenyl head group that extends into the II-III fenestration and an aliphatic tail positioned in the pore of Ca_v_3 channels (Zhao et al., 2019b; Huang et al., 2024). Hydrophobic contacts in the II-III fenestration are thought to enable allosteric, voltage-dependent modulation of Ca_v_3 channels (Zhao et al., 2019b; Huang et al., 2024), similar to how DHPs inhibit Ca_v_1 channels (Zhao et al., 2019a). Our structural analysis suggests that ML218 and Z944 do not bind to the II-III fenestration of Ca_v_1.4. Key residues involved in coordinating Z944 in Ca_v_3.1 (N592, F596, K1462) or ML218 in Ca_v_3.2 (N1003, F1007, K1503) are not conserved in Ca_v_1.4 (Zhao et al., 2019b; Huang et al., 2024), which could diminish the electrostatic and hydrophobic properties of the binding site (Supp. Fig.3). In our model of Ca_v_1.4, the favorable positioning of ML218 and Z944 in the III-IV fenestration (Fig.8) is intriguing given the structural similarities of these drugs and DHPs with dual activity on Ca_v_3 and Ca_v_1 subtypes (Supp. Fig.1). One such DHP is benidipine, which has an elongated structure like ML218 and Z944 (Supp. Fig.1B) and inhibits Ca_v_1.2, Ca_v_3.1, and Ca_v_3.2 in the low micromolar range (Furukawa et al., 1999; Furukawa et al., 2009). In the cryoEM structure of Ca_v_1.2, the phenylmethyl piperidinyl group of benidipine interacts with the hydrophobic pocket of the DHP binding site, which includes V1053, M1178, and M1509. These interactions may stabilize the binding of benidipine and contribute to its 10-fold higher potency compared to nifedipine (Wei et al., 2024). Considering that mutation of the corresponding residues in Ca_v_1.4 altered the IC_50_ of ML218 (Fig.9, Table 2), they may play a similar role in anchoring ML218 (Fig.8B,C). The importance of the hydrophobic interactions in the DHP site may also apply to DHP and DHP-like Ca_v_1 antagonists with large side chains such as (R)-GD10 and efonidipine (Supp.Fig.1B). In the low micromolar range, these drugs also block Ca_v_3 channels possibly through the III-IV hydrophobic pocket; polar residues, including that corresponding to T1007 in Ca_v_1.4, are not conserved in Ca_v_3 channels (Furukawa et al., 2009; Perez-Reyes et al., 2009; Gunduz et al., 2025). Thus, strong interactions of ligands with the hydrophobic DHP site may generally support Ca_v_3 and Ca_v_1 inhibition independent of the canonical threonine.

### Differential modulation of Ca_v_1.4 and Ca_v_1.2 by DHPs, ML218, and Z944

Among the Ca_v_1 subtypes, Ca_v_1.3 and Ca_v_1.4 are distinguished by their relatively low DHP potency which is ∼10-fold lower than for Ca_v_1.2 (Koschak et al., 2001; Xu and Lipscombe, 2001; Koschak et al., 2003). Efforts to uncover the molecular determinants in the DHP binding site that could underlie this difference focused on V1063 in rCa_v_1.2 (Fig.10). When mutated to the methionine residue present in hCa_v_1.3 (M1030, corresponding to M1004 in hCa_v_1.4), the IC_50_ of the mutant Ca_v_1.2 was slightly increased but did not match that of Ca_v_1.3 (Wang et al., 2018). Similarly, we did not find that the V1063M mutation in Ca_v_1.2 reproduced the ∼10-fold higher potency of ML218 observed for Ca_v_1.4 (Fig.10). Acidic amino acids upstream of the IIIS5 P1 region (EDD) in Ca_v_1.2 may allosterically couple DHP binding to non-conducting pore conformations (Wang et al., 2007). Structural modeling suggests that hydrophobic or neutral substitutions of these residues in Ca_v_1.3 together with M1030 may constrain the movement of the domain III pore helix upon DHP binding thus limiting pore closure (Wang et al., 2018). In our model of Ca_v_1.4, multiple poses of ML218 in the III-IV fenestration were energetically favorable (Fig.8, Suppl. Fig.3). ML218 could assume conformations that more strongly couple to pore closure in the context of Ca_v_1.4 than Ca_v_1.2. These conformations may not be accessible to Z944, leading to its lower potency for Ca_v_1.4 compared to ML218 (Table 2). The flexibility of ML218 in the Ca_v_1.4 binding site may also explain its ability to modestly shift the *V_h_* (Fig. 3C,D, Table 3). While nifedipine and the DHP agonist Bay K 8644 occupy the same binding site in Ca_v_1.2, a phenylalanine in IVS6 adopts a different conformation in the agonist vs antagonist bound state (Zhao et al., 2019a). Depending on the orientation of ML218, alterations in the positioning of this phenylalanine (F1430) and nearby M1426 could produce subtle shifts in S6 movements that underlie pore gating (Lenaeus et al., 2017; Yan et al., 2017).

### Implications for pharmacological analyses of Ca_v_ channels

Our study was motivated by our previous findings of modulatory effects of ML218 on Ca_v_1.4 channels in cone photoreceptors (Maddox et al., 2024). At a concentration of 5 μM, ML218 caused a 5-10% inhibition of Ca_v_1.4 *I_Ca_* and ∼2 mV hyperpolarizing shift in *V_h_*. While these effects did not reach statistical significance in mouse or ground squirrel cones, they are consistent with our analyses of Ca_v_1.4 in transfected HEK293T cells. While Z944 had only inhibitory effects on Ca_v_1.4 in our study (Fig.2), Davison and colleagues found that Z944 (5 μM) increased the amplitude of Ca_v_1.4 *I_Ca_* in mouse cones (Davison et al., 2022). Since splice variation can affect the pharmacological properties of Ca_v_ channels (Bourinet et al., 1999; Huang et al., 2013), responses to Z944 could differ for the Ca_v_1.4 splice variant(s) expressed in mouse cones and those used in our study.

Besides the retina, Ca_v_1.4 is also expressed in the pineal gland and at low levels in other tissues (Hemara-Wahanui et al., 2005; Schlick et al., 2010; Williams et al., 2022). Under some conditions, Ca_v_3 channels are upregulated in both the retina and pineal. For example, norepinephrine increases Ca_v_3.1 transcription in rat pinealocytes (Yu et al., 2015) and Ca_v_3.2 function is augmented in cone photoreceptors lacking functional Ca_v_1.4 channels (Maddox et al., 2024). In such cases, the use of Z944 or ML218 to pinpoint a contribution of Ca_v_3 channels might lead to erroneous conclusions. Studies employing systemic administration of these drugs to study behavioral or physiological endpoints should consider the potentially confounding effects of antagonizing retinal or pineal Ca_v_1.4 channels.

DHPs are among the most important classes of Ca_v_1 blockers and act mainly on Ca_v_1.2 channels in arterial smooth muscle in the treatment of vascular disorders such as hypertension and angina (Zamponi et al., 2015). A limitation of current DHP analogues is their weak ability to discriminate between Ca_v_1 subtypes. Elegant studies using knock-in mice expressing DHP-insensitive Ca_v_1.2 channels have highlighted the distinct physiological roles of Ca_v_1.2 and Ca_v_1.3 (Sinnegger-Brauns et al., 2004; Striessnig et al., 2006; Busquet et al., 2008). Ca_v_1.3 channels have emerged as potential drug targets in hyperaldosteronism, neuropathic pain, and Parkinson’s disease, and the development of Ca_v_1.3-selective compounds is a topic of intense research (reviewed in (Filippini et al., 2023)). Considering their similarities in DHP binding determinants, Ca_v_1.3 and Ca_v_1.4 may share the ability to be modulated by ML218 and Z944. The possibility that chemical modification of these drugs could lead to greater Ca_v_1-subtype specificity warrants further study.

