## Supplemental Figure 1 for "Inhibition of Ca_V_1.4 channels by Ca_V_3 channel antagonists ML218 and Z944"

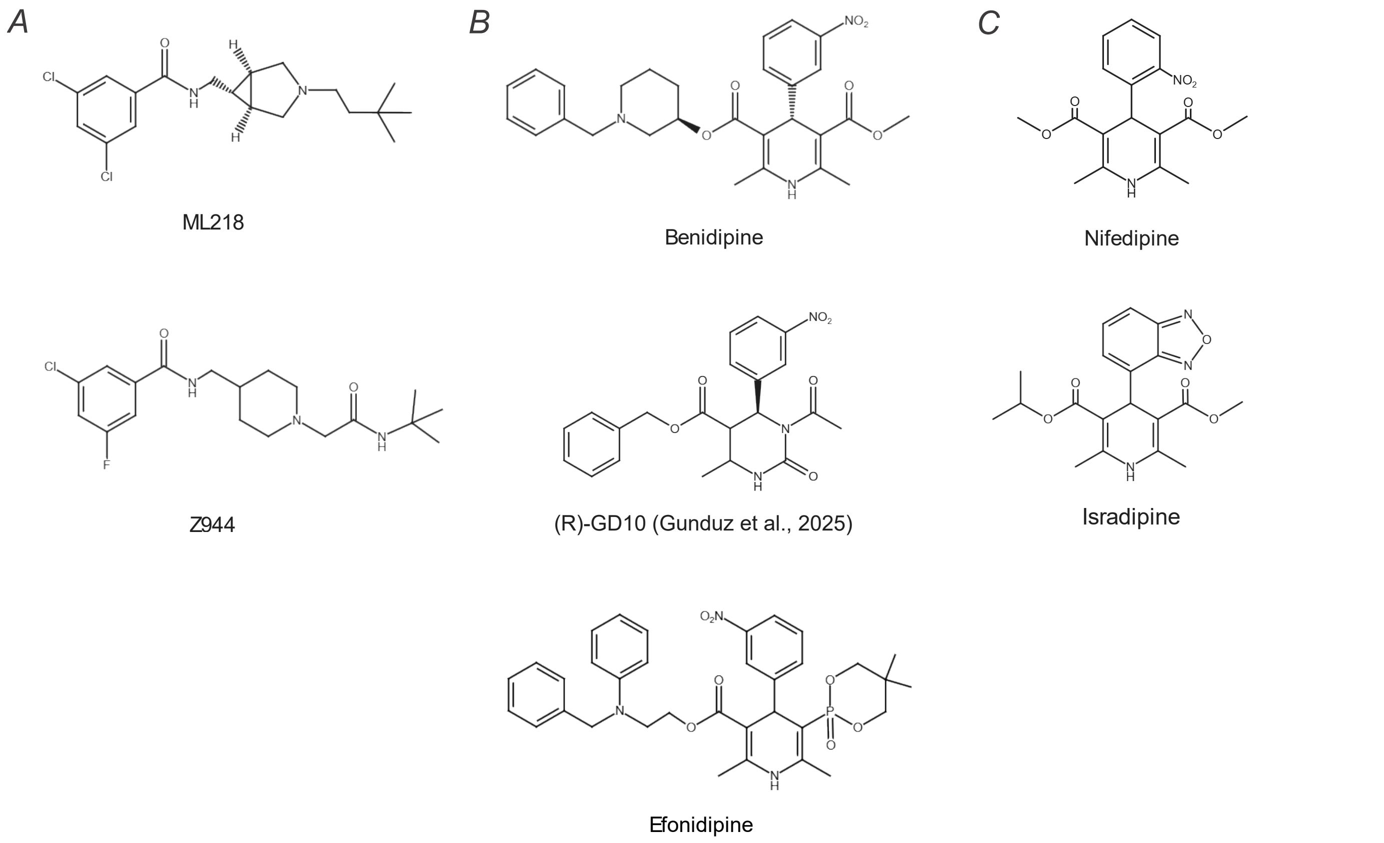


**Figure S1. Compounds with antagonist activity on Ca$_v$1 and/or Ca$_v$3 channels.** Unlike DHPs such as nifedipine and isradipine, the dual Ca$_v$1/Ca$_v$3 antagonists Benidipine, (R)-GD10, and Efonidipine have benzyl ester and phenylmethyl piperidinyl groups which form additional hydrophobic contacts in the III-IV fenestration of Ca$_v$1 channels {Zhao, 2019 #10411}{Wei, 2024 #10419}{Gunduz, 2025 #10417}, similar to the aliphatic tails of ML218 and Z944 in our study (Fig.8).
