## Supplemental Figure 2 for "Inhibition of Ca_V_1.4 channels by Ca_V_3 channel antagonists ML218 and Z944"

Supp Fig.2

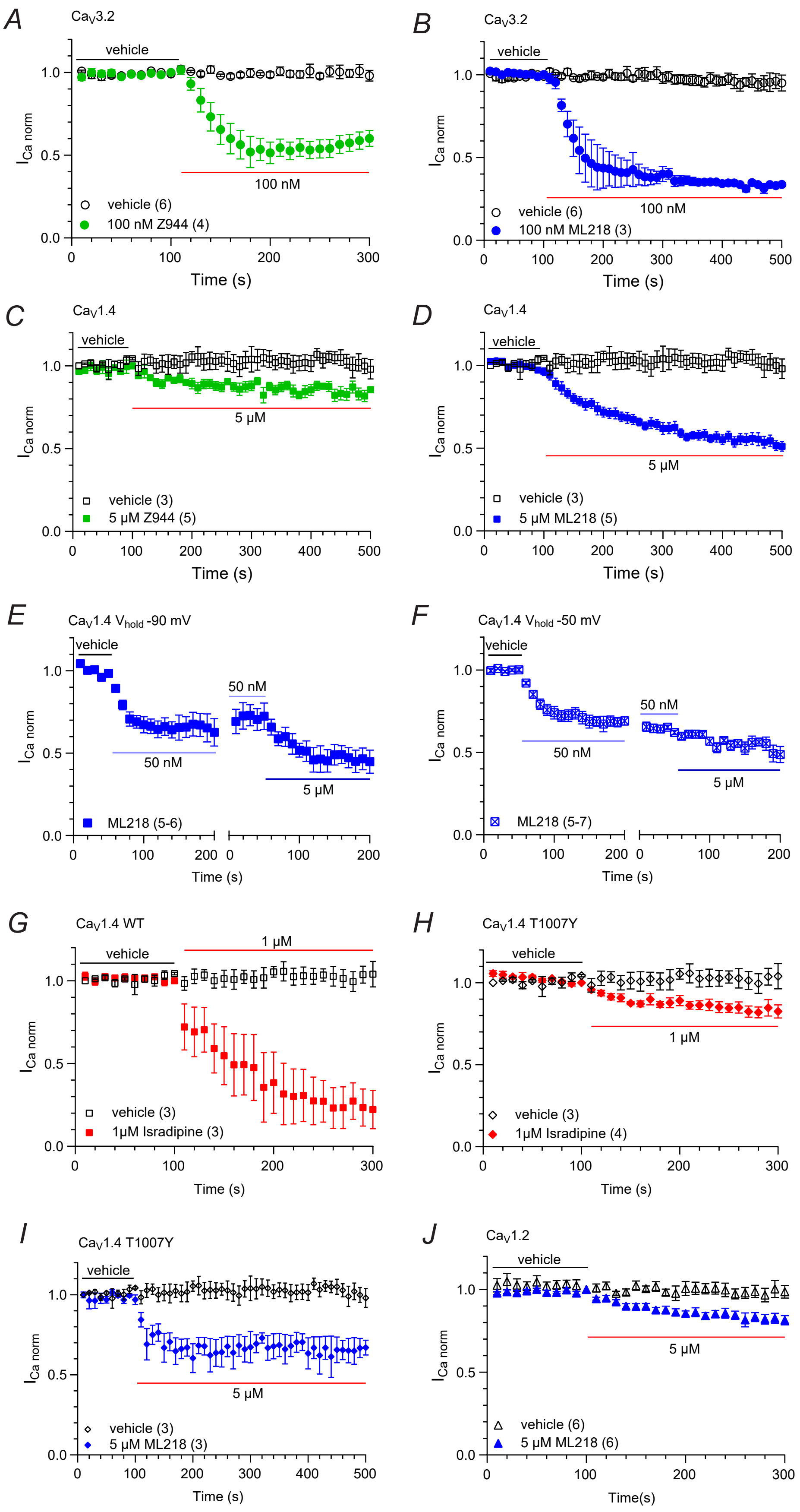


**Figure S2. Time course of drug responses. (A-J)** $I_{Ca}$ was evoked by 200-ms pulses from -90 mV (A-E, G-J) or -50 mV (F) to -20 mV (A,B) or 10 mV (C-J) at 0.1Hz. Peak currents were normalized to steady-state currents recorded with vehicle control. Points represent mean ± SEM. In E,F, gap indicates pause during which I-V relationships were measured. Parentheses indicate numbers of cells.
