## Supplemental Figure 3 for "Inhibition of Ca_V_1.4 channels by Ca_V_3 channel antagonists ML218 and Z944"

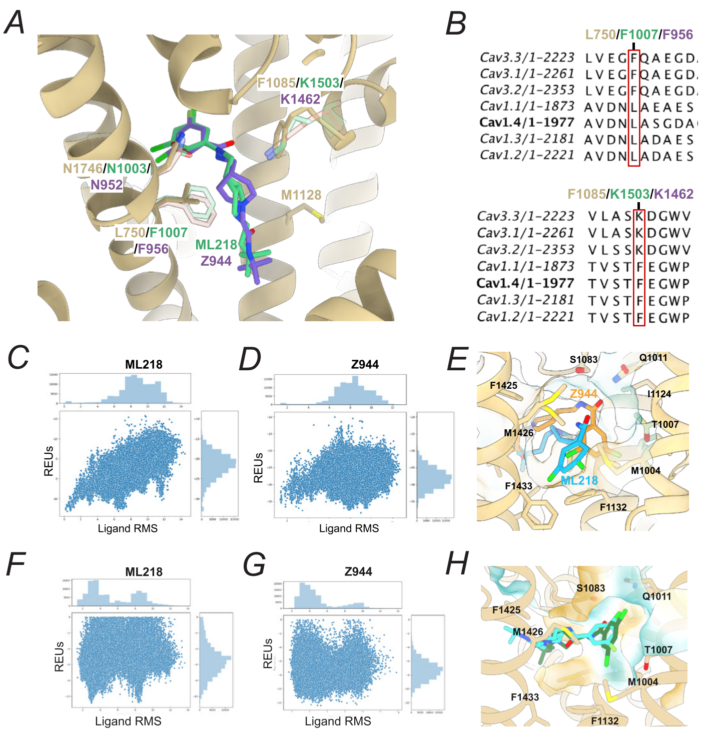


**Figure S3. Structural comparison and docking analysis of ML218 and Z944 in Ca$_V$1.4. (A)** Superposition of the experimental structures of Ca$_V$3.2–ML218 (PDB: 9ayk) and Ca$_V$3.1–Z944 (PDB: 6kzp) onto the Ca$_V$1.4 AF2 model. The full backbone of Ca$_V$1.4 is shown in beige, while only the local backbone fragments and interacting residues from Ca$_V$3.1 and Ca$_V$3.2 are displayed for clarity. K1503 in Ca$_V$3.1 and K1462 Ca$_V$3.2, which are important for ligand coordination, are replaced by F1085 in Ca$_V$1.4, altering the binding pocket electrostatics and hydrophobicity in that region. **(B)** Multiple sequence alignment of Ca$_V$3 and Ca$_V$1 channels. Boxed residues are major substitutions within the binding regions of ML218 and Z944. **(C)** Plot of ligand interface energy in Rosetta Energy Units (REUs) versus ligand RMSD for docked poses of ML218 in Ca$_V$1.4. A clear energy funnel is observed, with the lowest-energy pose (–33.19 REU) corresponding to the lowest ligand RMSD, indicating convergence. Additional local minima suggest that ML218 may adopt alternative low-energy configurations within the same binding region. **(D)** Plot of ligand interface energy (REU) versus ligand RMSD for Z944 docked into Ca$_V$1.4. A low RMSD pose aligns closely with the reference ML218 pose, but is not the global energy minimum. Nevertheless, this pose remains energetically favorable and supports the idea of a shared binding region. Alternative poses with lower interface energies were also observed. **(E)** Representative alternative docking poses for ML218 in Ca$_V$1.4 selected based on low interface energy. Despite geometric variation, these poses consistently form interactions with residues M1004, M1129, M1426, and Q1011, suggesting that these side chains contribute to defining a flexible yet structurally coherent binding environment. **(F)** Plot of the ligand interface energy (REU) versus ligand RMSD for ML218 with reference to the alternative pose in the DPH pocket two different low energy wells representing a 180° flip of the molecule, lowest REU value of -10.85 units. **(G)** Plot of the ligand interface energy (REU) versus ligand RMSD for Z944 lowest energy pose corresponds to -12.4 units. **(H)** Example poses for ML218 and Z944 according to Boltz2x prediction.
